# HES1 oscillations are required for cell cycle re-entry in oestrogen receptor positive breast cancer cells

**DOI:** 10.1101/2025.08.04.668440

**Authors:** Oliver Cottrell, Andrew Rowntree, Kunal Chopra, Eleanor Mackellar, Benjamin Noble, Hannah Dixon, Ciara Healy, Robert B. Clarke, Nancy Papalopulu

**Affiliations:** Division of Developmental Biology and Medicine, School of Medical Sciences, Faculty of Biology Medicine and Health, The University of Manchester, Michael Smith Building, Oxford Road, Manchester, M13 9PL, UK; Barts Cancer Institute, John Vane Science Centre, Charterhouse Square, Queen Mary University of London, London, EC1M 6BQ, UK; ApconiX, Alderley Park, Alderley Edge, SK10 4TG, UK; Manchester Breast Centre, Division of Cancer Sciences, School of Medical Sciences, Faculty of Biology Medicine and Health, The University of Manchester, Oglesby Cancer Research Building, Wilmslow Road, Manchester, M20 4GJ, UK

## Abstract

Long-term recurrence in breast cancer is driven by reactivation of dormant disseminated tumour cells (DTCs) and remains a major clinical challenge, particularly in oestrogen receptor positive (ER⁺) tumours. This process is underpinned by regulation of the cell cycle machinery that controls quiescence maintenance and exit. HES1, a Notch pathway transcription factor, regulates key cell cycle genes and has been shown to demonstrate protein expression oscillations.

Here, we sought to establish whether HES1 oscillations may regulate ER+ cancer cell quiescence and reactivation. To investigate this, we developed a fundamental *in vitro* model of cell cycle arrest and re-entry based on reversible CDK4/6 inhibition (CDK4/6i) with palbociclib, compatible with quantitative single-cell live-imaging of a knock-in endogenous HES1 reporter. Consistent with earlier findings, HES1 exhibited ∼24-hour protein oscillations in cycling cells demonstrating a reproducible dip in protein expression prior to S-Phase. During CDK4/6i-mediated arrest, the ∼24h HES1 oscillation was lost, HES1 levels were maintained at a moderately higher level and HES1 exhibited smaller dips. Similar changes were observed in unperturbed, spontaneously quiescent cells. Following release from CDK4/6i and cell cycle re-entry, these alterations were reversed and the characteristic G1/S HES1 dip was observed. Preventing this dip at the point of release, by inducibly sustaining HES1 with a Tet-On system, upregulated the cell cycle inhibitor p21, impeded cell cycle re-entry and induced cell death. These findings suggest that manipulating HES1 dynamics could represent a promising therapeutic approach to prevent reactivation of dormant tumour cells.

**Significance Statement:** Breast cancer can recur years after initial treatment due to reactivation of dormant tumour cells. Understanding how these cells exit dormancy is crucial for preventing relapse. We investigated HES1, a transcription factor with rhythmic protein oscillations, and its role in regulating quiescence in oestrogen receptor-positive (ER⁺) breast cancer cells. Using live-cell imaging and a reversible cell cycle arrest model, we show that HES1 dynamics change during dormancy and reactivation, and that disrupting these oscillations prevents cell cycle re-entry and induces cell death. These findings reveal HES1 protein dynamics as a potential therapeutic vulnerability and highlight a novel strategy to target dormant cancer cells to prevent their reactivation.

## Introduction

Breast cancer is the leading class of female cancer worldwide, with around 1 in 7 women now expected to develop the cancer in their lifetime [1–2]. In recent decades, mortality rates have been markedly reduced by earlier diagnosis and improved targeted therapeutic options. Despite advances in 5-year survival however, the risk of relapse, metastasis and fatality persists beyond that point. This is particularly evident in ER+ luminal breast cancers, where the rate of recurrence within 20 years ranges from 15-38% [3].

Indeed, long-term distant recurrence, can present up to 20 years following the termination of primary adjuvant therapy in ER+ breast cancer patients [3–4]. This points towards long periods of clinical dormancy, where tumours remain in stasis at secondary sites, retaining the propensity to later resume growth and form overt metastases. This can occur at the level of the tumour mass, where the proliferation rate of micro-metastases is balanced by immune-surveillance and limited vascularisation, as well as at the single-cell level, where individual DTCs are functionally quiescent [5–6].

Recent research has significantly advanced our understanding of DTC dormancy. Rather than arising from reproducible mutations, dormancy has been associated with epigenetic reprogramming, with repressive chromatin marks shaping transcriptional programmes that sustain quiescence [7–8]. These transcriptional programmes are not static but can be shaped or re-wired by cues from the tumour microenvironment (TME), where immune cells, stromal cells, and hypoxia have emerged as key modulators of dormancy and reactivation [9–14]. Despite these insights, no therapeutic options for managing dormant DTCs currently exist, highlighting the urgent need to better understand the mechanisms and pathways by which DTCs transition between dormancy and reactivation.

The Notch signalling pathway is a potent regulator of cell fate and proliferation during development across multiple tissues, including the mammary gland [15–16]. Dysregulation of Notch signalling has been implicated in tumorigenesis across various contexts [17–19]. Moreover, Notch signalling has been shown to be important for engraftment of dormant niches in the bone marrow by breast DTCs [20], while Notch signalling is strongly upregulated in NR2F1 agonist-induced dormancy [21].

Downstream of Notch signalling, HES1 a transcriptional repressor, has also been implicated in cancer. HES1 is upregulated in breast cancer overall, as well as luminal A breast cancer specifically, compared to normal mammary tissue [22–23] (Fig. S1A-B). Furthermore, in hormone therapy resistant ER+ breast cancer cell lines and patient-derived xenograft (PDX) tumours, HES1 is upregulated in the cancer stem cells which are likely to be the source of breast cancer recurrence [24]. At the molecular level, HES1 modulates proliferation by transcriptional repression of several elements of the cell cycle machinery, including CycD1, CycE2, CycA2, E2F, p21 and p27 [25–29]. Thus, overexpression of HES1 in an ER+ breast cancer cell line, reduced proliferation by repression of E2F-1 [26], while HES1 downregulation by HES6 had the reverse effect [30].

These studies have illustrated the relevance of HES1 to breast cancer with an emphasis on the mean levels of HES1 expression. However, developmental studies have indicated that Notch pathway components regulate cell fate and proliferation by virtue of their dynamic gene expression patterns. Specifically, Notch ligands (Dll1), direct Notch targets (HES1, HES5) and their downstream targets (ASCL1, NGN2, NGN3) have been shown to periodically fluctuate (i.e. oscillate) at the protein level. The characteristics of these oscillations (amplitude, frequency and phase), rather than mean expression levels alone, have been shown to functionally regulate somitic, neural and pancreas development [31–35]. Moreover, manipulating HES1 from wild-type oscillatory dynamics to sustained expression in adult neural stem cells (NSCs) has been shown to reduce proliferation rate [36–37] and prevent re-entry from cell cycle arrest [38], via differential regulation of p21 [37–38].

Motivated by the functional importance of dynamics, our recent work investigated HES1 oscillations in ER+ breast cancer [39], uncovering a cell cycle-associated periodicity of ∼25h, which was similar to the average cell cycle length (∼24h). An oscillatory dip in HES1 expression reliably occurred prior to S-phase, ∼10-14 hours prior to the next division, leading us to hypothesise that HES1 oscillations are required for the G1/S transition.

Here, we investigate the relevance of HES1 oscillations specifically to cell cycle re-entry from quiescence. To study tumour dormancy and recurrence with multi-day quantitative single-cell live-imaging with high temporal resolution, an in vitro model of cell cycle arrest and re-entry was developed using reversible CDK4/6 inhibition (CDK4/6i) with palbocilcib in an ER+ breast cancer cell line. We show that during arrest the cell cycle-associated ∼24h HES1 oscillations are absent but are rescued upon cell cycle re-entry. Manipulating HES1 oscillations, using an inducible expression system, impeded cell cycle re-entry, induced cell death and upregulated the cell cycle inhibitor p21. This indicates a bi-directional relationship whereby HES1 is regulated by, but also regulates, the cell cycle. Together, our findings suggest that unperturbed HES1 dynamics are required for successful re-entry from cell cycle arrest and that interfering with HES1 dynamics may have therapeutic relevance towards managing dormant tumour cells.

## Results

### Modelling reversible cell cycle arrest with CDK4/6 inhibition

First, we established an in vitro model of ER⁺ breast cancer dormancy compatible with multi- day, single-cell imaging of HES1 protein dynamics. Due to the capacity for CDK4/6 inhibitors to invoke reversible G0/G1 arrest *in vitro* [40–41], we reasoned that they could be used to mimic the fundamental cell cycle features of entry into and exit from arrest observed during dormancy and reactivation. Thus, we treated ER+ MCF-7 cells with the CDK4/6 inhibitor, palbociclib, which markedly suppressed population growth, Ki67 positivity (stains late G1–M cells; [42]), and EdU incorporation (marks S-phase; Fig. 1A–D), indicative of robust cell cycle arrest. Upon release from palbociclib, population growth, Ki67 staining, and EdU incorporation recovered within one day of drug withdrawal (Fig. 1A–D), signifying a high degree of reversibility.

**Figure 1.**
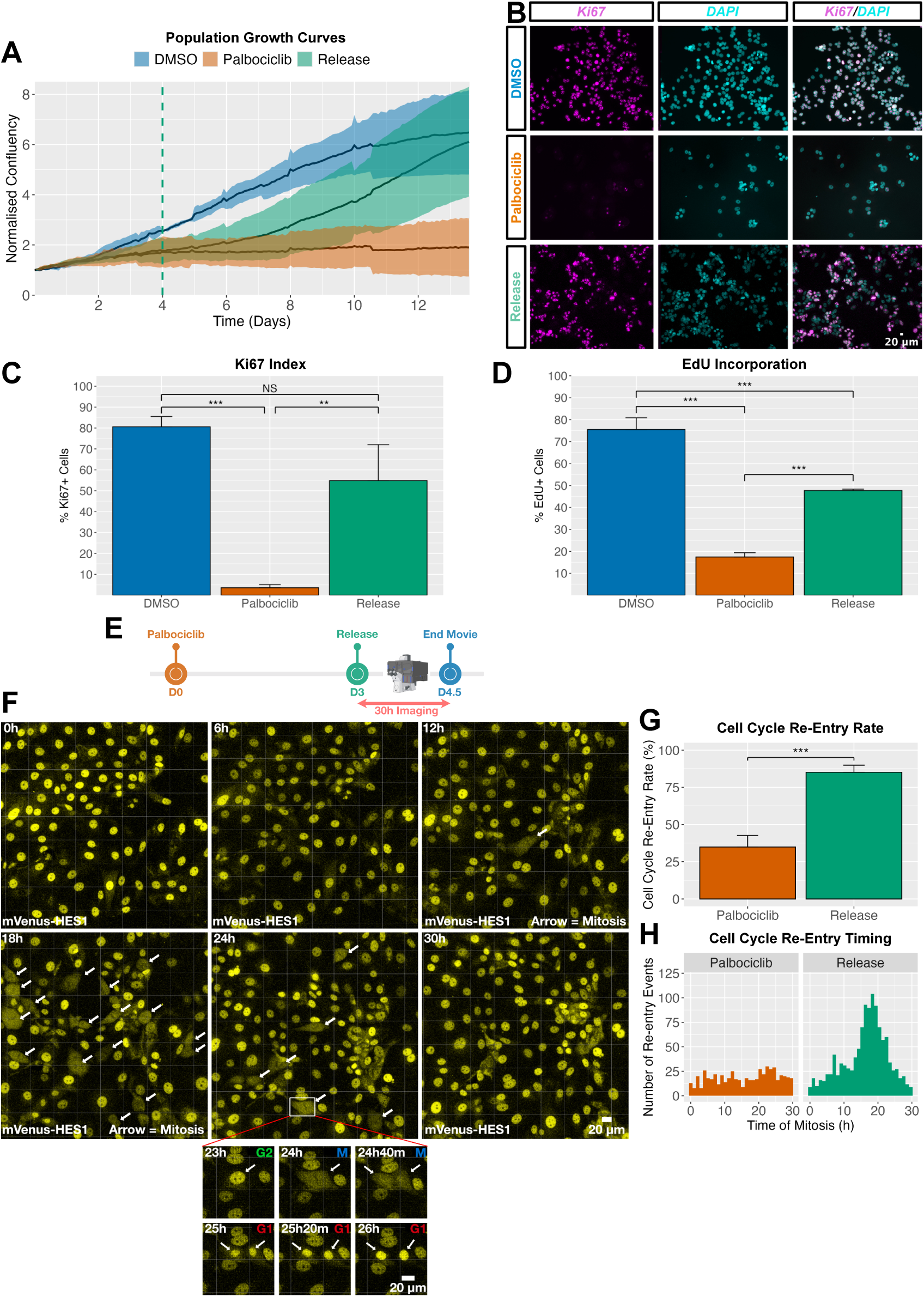
Palbociclib-induced reversible G0/G1 arrest in MCF-7 cells to model ER+ breast cancer dormancy and re-awakening. (A) Population-level growth curves of MCF-7 cells treated with DMSO (proliferative control), continuous palbociclib, or palbociclib followed by release into proliferative media after 4 days. Lines represent mean of *N* = 4 biological replicates; shaded areas show SD. Confluency was normalised to baseline. Green line indicates time of release. (B) Representative immunofluorescence images of cells stained for the proliferation marker Ki67. Cells were released from palbociclib after 3 days herein. (C) Automated quantification of Ki67⁺ cells from (B). Bars represent mean ± SD from *N* = 3 biological replicates, *n* = ∼1000 cells per group. Independent *t*-test: \*\*\**p < 0.001, **p < 0.01*. (D) Fraction of EdU⁺ cells after a 24 h pulse, measured by flow cytometry. Bars represent mean ± SD from *N* = 3 biological replicates (*n* = 11,929 DMSO; 9,014 palbociclib; 10,626 release). Independent *t*-test: \*\**p* < 0.001. Higher proliferation is inferred here than (C) due to fresh medium being supplied during the EdU chase, providing new serum and transient proliferative stimulus (see SI Methods). (E) Experimental timeline (F-H). mV-HES1 cells were released from palbociclib after 3 days and imaged for 30h to measure mitosis frequency. (F) Representative time-lapse snapshots from live-imaging of released cells. Imaging commenced ∼1h post-release. Endogenous mVenus-HES1 signal was used as a nuclear marker. Arrows indicate mitotic events. Inset shows a cell undergoing division. (G) Percentage of cells in (F) undergoing mitosis within 30h post-release, defined as ‘re-entry’ events. Bars show mean ± SD from *N* = 3 replicates (*n* = 560 palbociclib; 1,162 release cells). Independent *t*-test: ***p < 0.001. (H) Histogram showing the timing of re-entry events in (F), pooled across *N* = 3 replicates.

Live-imaging of released cells using a HES1 reporter as a nuclear marker (Fig. 1E; reporter shown in Fig. 2A) revealed that released cells proceeded largely synchronously from G0/G1 arrest to division, as previously reported in other cell lines [40], reaching mitosis most commonly ∼18-19h following palbociclib removal (Fig. 1E-G) such that 85% of cells underwent mitoses within 30 hours of drug withdrawal (Fig. 1F). This synchronicity was quickly lost over subsequent generations in the released cells.

**Figure 2.**
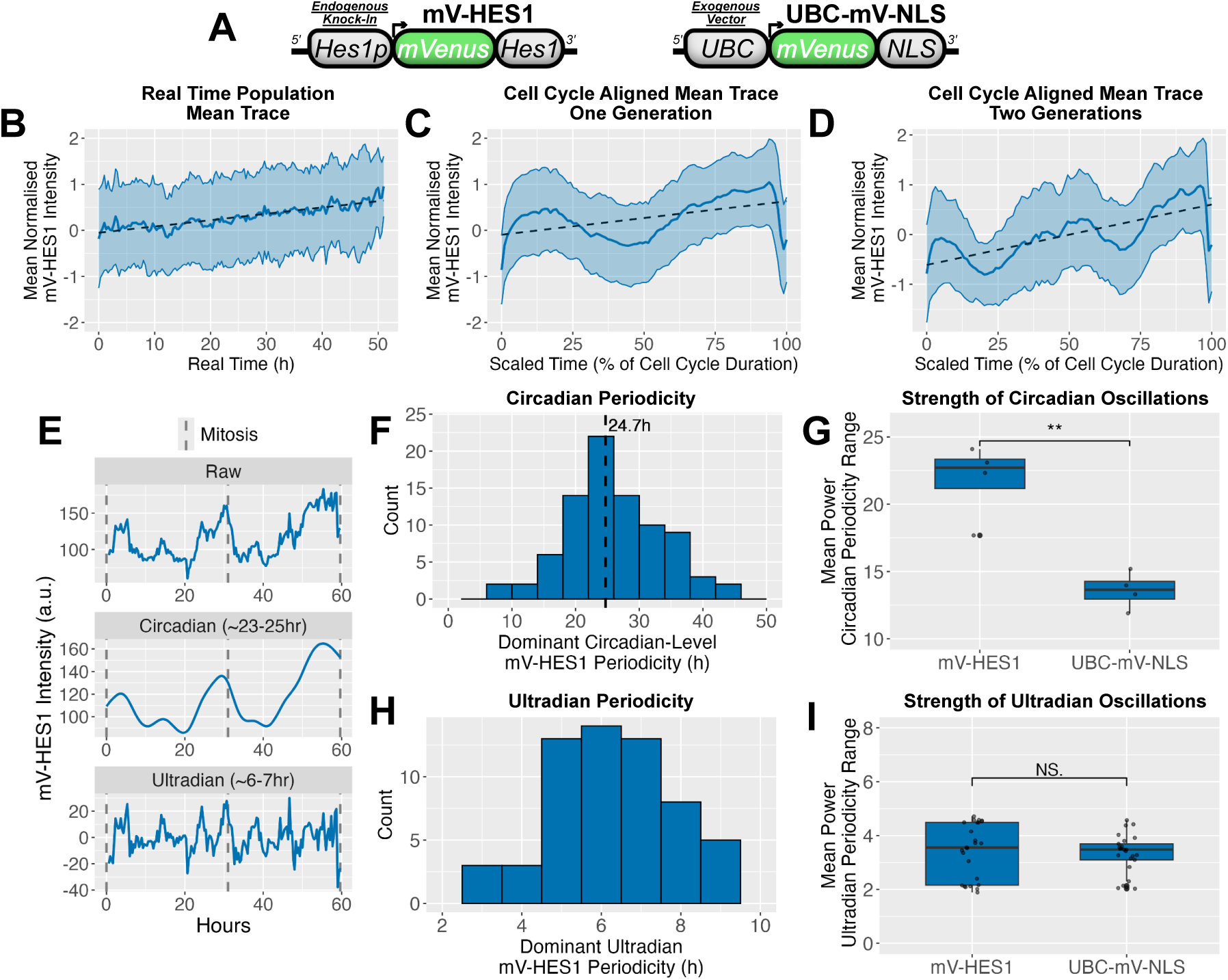
HES1 exhibits single-cell oscillations with circadian-level frequency in proliferative MCF7s. (A) Schematic of the endogenous mVenus-HES1 fusion reporter in MCF-7 cells (left), alongside the exogenous UBC-mVenus-NLS control reporter (right). (B) Single mV-HES1 cells were tracked from mitosis to mitosis, then individual time-series were Z-score normalised and aggregated to generate a population mean trace. Averaging across real-time points revealed no coherent population-level oscillations. *N* = 3 independent experiments *; n* = 90 cells. (C) As in (B), but time-series were pseudo-synchronised by scaling time as a percentage of each cell’s cell cycle duration, revealing oscillations associated with cell cycle progression. (D) As in (C), but showing two consecutive cell cycle generations (mother–daughter traces). (E) Representative single-cell mV-HES1 time-series. Raw signal (top) is shown alongside detrended signals isolating circadian-level (middle) and ultradian (bottom) oscillations. (F) Histogram of circadian-level periodicities estimated by Lomb–Scargle periodogram (LSP). *N* = 3 experiments; *n* = 90 cells. (G) Comparison of oscillation power within the circadian frequency range (20-30h) between proliferative mV-HES1 and UBC-mVenus-NLS signal, based on LSP analysis (F). Each point represents mean power at a given circadian-level frequency averaged across cells and replicates. Independent t-test: **p < 0.01. (H, I) As in (F) and (G) but for ultradian oscillations (4-8h range) using appropriately detrended time-series.

It remained unclear whether palbociclib treatment produced a heterogenous population in which a refractory subset continued to proliferate while others were arrested, or whether all cells experienced comparable cell cycle arrest. To distinguish between these possibilities, we quantified the fraction of cells that divided within a 100h window, from day 3 to day 7 of palbociclib treatment. Approximately 75% of control cells divided at least once and 50% divided twice within this window which fell to 25% and 1.5% respectively with palbociclib treatment (Fig. S2E-F). Moreover, the number of dividing cells gradually declined over the time-course before ceasing entirely (Fig. S2G). Together, these data indicate that while a small fraction of cells may continue to divide transiently, the vast majority ultimately enter arrest, arguing against the presence of a stable, refractory proliferative subpopulation.

Finally, a small reduction in cell viability from 93% to 81% was observed upon drug treatment (Fig. S2A-D), which has previously been observed [43–44], however this small proportion of death (12%; apoptosis plus necrosis, Fig S2D) is unlikely to be sufficient to explain the overall population-level growth arrest (Fig. 1A), framing decreased proliferation as the predominant effect. Together, these data indicate that palbociclib can be used to readily induce reversible G0/1 arrest in MCF-7 cells, serving as an *in vitro* model of experimental dormancy and reactivation, in a manner that is amenable to single-cell live-imaging.

### HES1 exhibits 24h oscillations which are correlated with cell cycle progression

To observe HES1 protein dynamics in cycling cells, we used MCF-7 cells expressing an endogenous HES1 fusion reporter (mV-HES1) previously generated by CRISPR-mediated tagging [39], enabling real-time single-cell imaging of HES1 protein levels (Fig. 2A). Cells expressing a constitutively driven nuclear mVenus reporter (UBC-mV-NLS) served as a non-oscillatory control.

Proliferative cells exhibited HES1 oscillations at the single-cell level, which were asynchronous across the population and could only be observed when single cell traces were aligned from mitosis to mitosis (Fig. 2B-D). In most cells, HES1 expression was high at the beginning of a cell cycle, then temporarily declined (herein termed the “dip”), before accumulating again, reaching a second high level prior to mitosis (Fig. 2C-E). Viewed over consecutive cell divisions (Fig. 2D-E), this cell cycle-correlated oscillation was found to have a median periodicity of 23- 25h using Lomb-Scargle Periodogram (LSP) (Fig. 2F) and Autocorrelation Function (ACF) analysis pipelines (Fig. S4). The term ‘circadian-level’ is used to describe this circa 24h periodicity herein, without implying a specific causative connection to the circadian clock. This circadian-level periodicity was significantly more powerful than non-oscillatory control UBC- mV-NLS time-series (Fig. 2G; Fig. S3) and matched the average length of the cell cycle in these cells [39].

In line with our earlier report [39], shorter ultradian-level periodicity was also contained within HES1 time series from proliferative cells, which was apparent when the traces were detrended to remove the longer periodicity (Fig. 2E). These nested ultradian oscillations demonstrated a median periodicity of 6-7h (Fig. 2H), however their power was not significantly different from UBC-mV-NLS controls (Fig. 2I), indicating that ultradian HES1 fluctuations are weak or indistinguishable from noise in proliferative cells.

### HES1 expression dip coincides with the G1/S transition

We previously showed that HES1 exhibits ∼24 h oscillations that are phase-locked to cell cycle progression in most cells, with a reproducible dip occurring 10–14h prior to mitosis and coinciding with the G1/S transition (Fig. 3A–B) [39]. This relationship was initially established using a live PCNA reporter to mark S-phase entry. Here, we extended this analysis by co-imaging HES1 with established G1/S regulators, CDK2 and p21.

**Figure 3.**
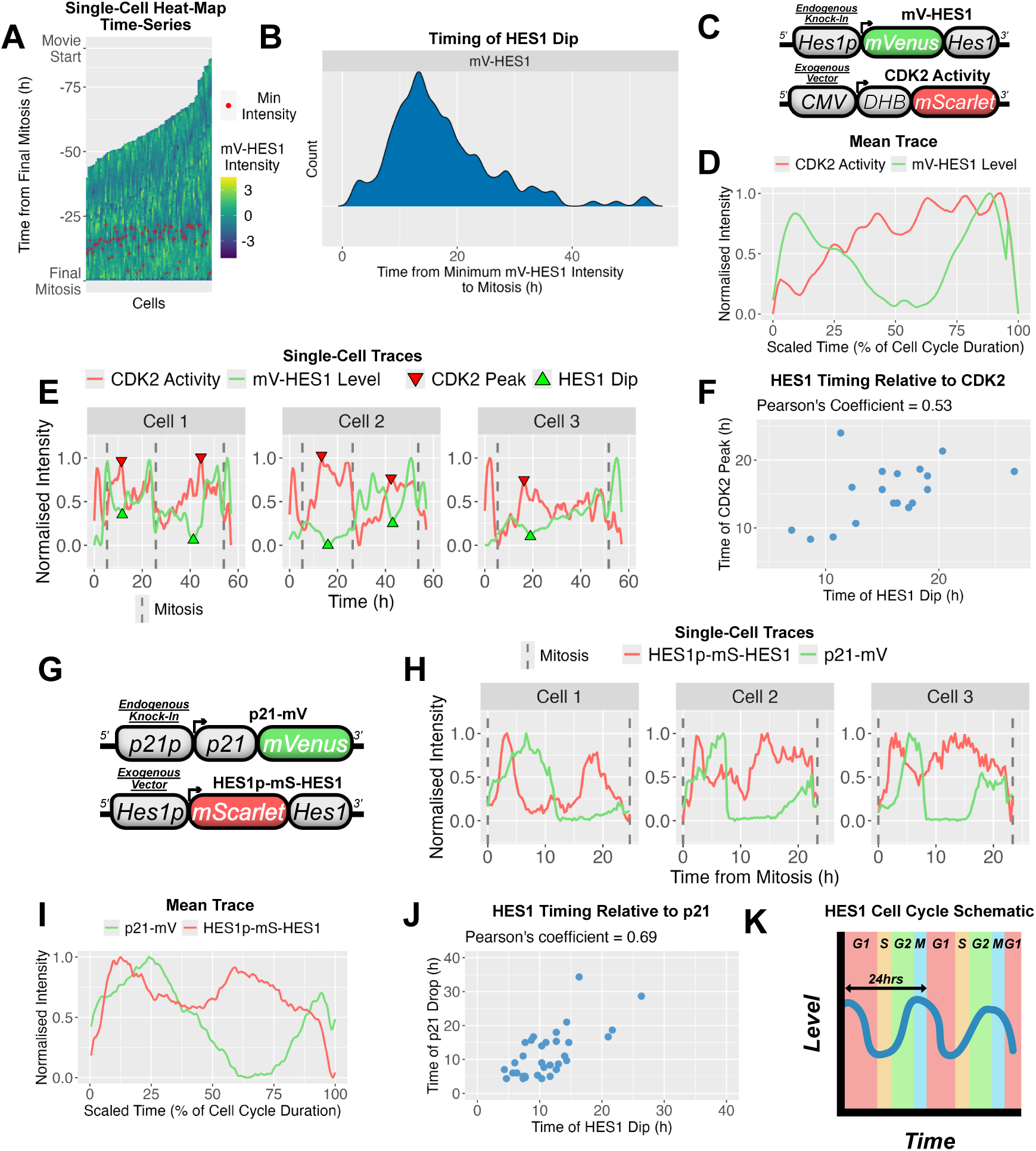
HES1 circadian-level oscillations correlate with cell cycle phase with the HES1 dip coinciding with G1/S. (A) Z-score normalised circadian-level mV-HES1 time-series from proliferative cells displayed as heatmaps. Each x-axis point represents a distinct cell; each trace includes two consecutive cell cycle generations with three mitoses. Traces are aligned (T = 0h) to the final. Red dots indicate the time of minimum mV-HES1 intensity in the daughter generation. *N* = 3 experiments, *n* = 90 cells. (B) Density plot showing the timing of minimum HES1 intensity from (A), relative to final mitosis (T = 0h). (C) Schematic of constructs used in (D-F). DHB-based vector was incorporated into mV-HES1 cells to visualise CDK2 activity alongside HES1 expression. (D) Mean trace of time-series from dual-reporter live-imaging of mV-HES1 and DHB-mScarlet (CDK2 activity) over a single cell cycle. Traces were normalised to their minimum and maximum values to visualise relative dynamics. *N* = 2 experiments; *n* = 14 cells. (E) Representative single-cell time-series from (D). (F) Scatter plot showing the timing of the HES1 dip vs the CDK2 maximal activity (G1/S) in single cell from (D, E). (G) Schematic of the endogenous p21-mV fusion-reporter and exogenous HES1p-mS-HES1 reporter cell line used in (H-J). (H) Representative single-cell time-series from dual-reporter live-imaging of p21-mV and HES1p-mS-HES1 over one cell cycle. Normalised as in (D). (I) Mean trace of time-series from (H). *N* = 3 experiments; *n* = 41 cells. (J) Scatter plot showing the timing of the HES1 peak vs p21 decline (G1/S) in single cells from (H, I). (K) Schematic illustrating the alignment of the ∼24h HES1 oscillation with discrete cell cycle stages.

Dual-colour live imaging of HES1 and a CDK2 activity reporter revealed an inverse relationship, with the HES1 dip aligning with peak CDK2 activity (Fig. 3C–F). Because rising CDK2 activity marks commitment to S-phase [45–46], this correlation supports the HES1 dip as occurring with precise timing relative to the G1/S transition. Similarly, co-imaging HES1 with p21 showed that both proteins oscillate across the cell cycle with a delayed phase relationship (Fig. 3G–I). The timing of the HES1 dip was well correlated with the timing of the p21 decline (Fig. 3J), a hallmark of the G1/S transition [47–48], with the HES1 dip typically preceding the p21 drop (Fig. 3I). This further supports the placement of the HES1 dip at, or shortly prior to, G1/S. Together, these co-imaging analyses demonstrate that the HES1 dip is tightly and reproducibly aligned with molecular markers of G1/S, supporting the conclusion that the ∼24h HES1 oscillation is structured by cell cycle progression (Fig. 3K).

### HES1 24h protein expression oscillations are absent in arrested cells

We next investigated how HES1 protein dynamics are altered during cell cycle arrest by quantitative live imaging of palbociclib-treated and vehicle-treated (DMSO) mV-HES1 cells (Fig. 4A). The ∼24 h periodicity in HES1 expression was markedly reduced in palbociclib-arrested cells, evident in raw traces (Fig. 4A) and confirmed by Lomb–Scargle periodogram (LSP) analysis (Fig. 4B–C), indicating that circadian-level HES1 oscillations are lost during arrest. No corresponding change was observed in UBC-mV-NLS control cells treated with palbociclib (Fig. 4C; Fig. S3).

**Figure 4.**
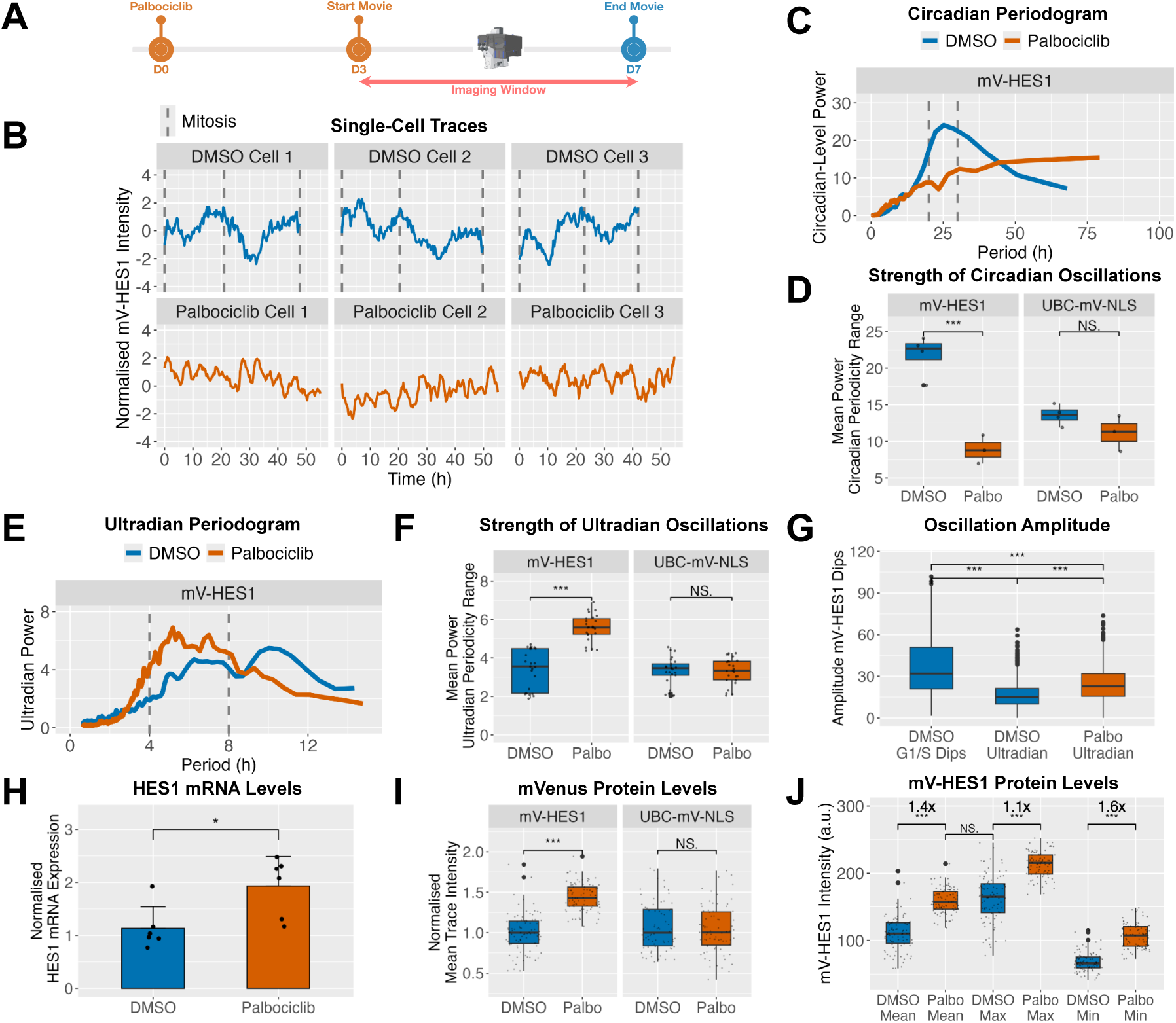
Circadian-level HES1 oscillations are absent during arrest, while levels are moderately elevated but remain within the physiological range. (A) Experimental timeline. mV-HES1 cells were treated with palbociclib or DMSO for 3 days and then continuously live-imaged for 4 further days under the same conditions. (B) Representative Z-score normalized time-series of mV-HES1 cells from each condition. (C) Mean normalised LSP power spectra from detrended time-series data in (B), showing oscillatory power across periodicities. Grey lines indicate circadian range used for quantification in (D). *N* = 3 independent experiments; *n* = 90 cells. (D) Comparison of circadian range oscillatory power (20-30h) between mV-HES1 and UBC- mVenus-NLS cells ± palbociclib, based on data in (C). Each point represents mean power at a given circadian-level frequency averaged across cells and replicates. ANOVA with Tukey’s post- hoc test. ***p < 0.001. (E, F) As in (C) and (D), but for ultradian oscillations (4-8h range). (G) Amplitude of every peak–dip pairs in various mV-HES1 traces, compared across circadian- and ultradian-detrended DMSO traces and ultradian-detrended palbociclib traces. *N* = 3 experiments; *n* = 90 cells per condition. ANOVA with Tukey’s post hoc test: \*\**p* < 0.001. (H) HES1 mRNA levels measured by RT–qPCR after 3 days of palbociclib treatment, expressed as fold-change relative to DMSO. Bars show mean ± SD from *N* = 6 experiments. Independent *t*-test: *p* < 0.05. (I) Mean mVenus fluorescence intensity per cell, calculated from raw time-series and normalised to the median DMSO value for each cell line. *N* = 3 experiments; *n* = 90 cells per condition. Independent *t*-test: \*\**p* < 0.001. (J) As in (I), showing mean, maximum, and minimum intensity per mV-HES1 cells ± palbociclib. Annotated fold-changes indicate ratios of palbociclib to DMSO medians. *N* = 3 experiments; *n* = 90 cells per condition. Independent *t*-test: \*\**p* < 0.001.

In contrast, the relative power of HES1 ultradian oscillations was increased in arrested cells compared to both proliferative mV-HES1 cells and UBC-mV-NLS controls (Fig. 4D–E). Although the amplitude of ultradian dips was enhanced, these remained smaller than the G1/S-associated dips observed in proliferative cells (Fig. 4F), indicating reduced overall HES1 variability during arrest. The observations that HES1 ultradian oscillations in proliferative cells are sub-detectable, exhibit reduced amplitude and do not correlate with cell cycle phase, suggest they are unlikely to be important in the context of normal cell cycle progression. Thus we focussed on the interaction between circadian HES1 oscillations and the cell cycle herein.

To determine whether these changes were specific to CDK4/6 inhibition, we analysed spontaneously non-dividing (SND) mV-HES1 cells in proliferative cultures (Fig. S5A–B). SND cells exhibited the same loss of circadian-level oscillations and relative enhancement of ultradian periodicity (Fig. S5C–F), indicating that these features are a general consequence of cell cycle arrest rather than a drug-specific effect.

Similarly, we investigated the small minority of cells (1.5%; Fig. S2F) which successfully completed a full cell cycle during palbociclib treatment, to determine whether they resembled normal cycling cells. Most palbociclib dividers demonstrated elongated cell cycles with a median duration of 46.3h compared to 23.3h in vehicle-treated dividers (Fig. S6B), which weakened the circadian character of the HES1 cell cycle dynamics over the population (Fig. S6C-D). Ultradian periodicity was stronger than in control proliferative cells (Fig. S6E-F). Among the palbociclib dividers, those with shortest cell cycles (see Cell 1-2; Fig. S6A) best resembled standard proliferative HES1 dynamics, by exhibiting clear biphasic expression with two dominant peaks, separated by the dip-phase. These data suggest that HES1 dynamics sensitively reflect the degree to which the cell cycle is progressing normally.

### Conservation of cell cycle-related HES1 dynamics in other breast cancer models

To assess whether the HES1 dynamic behaviours observed in MCF-7 cells extend beyond ER⁺ breast cancer, we engineered an endogenous mScarlet-HES1 fusion reporter in SUM149 cells, a triple-negative breast cancer line (Fig. S7A-C). Correct in-frame knock-in was confirmed by genotyping and sequencing (Fig. S7C-D), while snapshot immunofluorescence analysis demonstrated that HES1 expression heterogeneity was preserved relative to parental cells (Fig. S7E), indicating faithful reproduction of endogenous dynamics.

Similar to MCF-7s, proliferative SUM149 cells displayed cell-cycle-associated HES1 oscillations (Fig. S7F; Fig. S8A-B) with a dominant ∼26h periodicity that matched their median cell cycle duration (Fig. S8C-D), although the normalised circadian-range power was lower than in MCF- 7s (Fig. S8E). To test whether these oscillations were similarly sensitive to cell cycle arrest, SUM149 cells were treated with palbociclib at a 10-fold higher concentration than required for MCF-7s, consistent with prior reports [49]. Efficient population-level arrest was confirmed by longitudinal growth analysis (Fig. S8F). As in MCF-7s, palbociclib-arrested SUM149s lacked a consensus peak in the circadian-periodicity range, and power in this range was significantly reduced compared to proliferative counterparts (Fig. S8H-I). However, unlike MCF-7s, ultradian periodicity was diminished rather than enhanced, indicating a degree of cell-line variability for this frequency (Fig. S8J-K).

Together, these data demonstrate that circadian-level, cell cycle-associated HES1 oscillations are a conserved feature of proliferative breast cancer cells and are consistently lost upon cell cycle arrest across distinct molecular cell line models.

### HES1 is maintained at a higher level during arrest but remains with the physiological range

In addition to modulating HES1 dynamics, palbociclib treatment increased HES1 expression, with a 1.9-fold rise in mRNA levels by RT-qPCR (Fig. 4G) and a 1.4-fold increase in mean protein levels based on single-cell live imaging of mV-HES1 cells (Fig. 4H). This effect was specific to HES1, as no change was observed in control UBC-mV-NLS cells (Fig. 4H). Moreover, HES1 protein was also upregulated 1.4-fold in SND cells compared to proliferative counterparts (Fig. S5G), indicating this is not specific to palbociclib treatment but extends to unperturbed models of arrest as well. Notably, while palbociclib raised mean HES1 protein levels, these remained within the normal dynamic range observed during cycling. Mean levels in arrested cells were not significantly different from the maximum levels seen in proliferating cells (Fig. 4I). The increase in maximum HES1 intensity during arrest was modest (1.1-fold), whereas the minimum intensity rose more substantially (1.6-fold) compared to controls. These data indicate that palbociclib elevates the baseline, rather than the maximum, HES1 expression. Thus, palbociclib maintains HES1 at the highest levels normally seen in proliferating cells, preventing their decline rather than increasing it beyond the physiological range.

Together, the data indicates that HES1 oscillatory dynamics are profoundly altered during cell cycle arrest, manifested primarily as a loss of the circadian-level, cell cycle-associated, oscillations. Additionally, HES1 expression is maintained at a moderately elevated level (1.4- fold) and exhibits less pronounced dips compared to cycling cells.

### HES1 24h oscillations return upon cell cycle re-entry

To determine whether circadian-level HES1 oscillations could be rescued by cell cycle re-entry, arrested mV-HES1 cells were released from palbociclib into growth media and monitored by single-cell live imaging beginning ∼1 h later (Fig. 5A). A PCNA-based reporter (Fig. 5B), which becomes punctate in S-phase [50], was used to visualise the G1/S transition following release (Fig. 5C). Circadian-level oscillations returned in released cells, exhibiting characteristic cell cycle–associated HES1 dynamics, with peaks at mitoses, interspaced by G1/S dips (Fig. 5D). Periodicity analysis (ACF and LSP) revealed a dominant period of 22–26h, similar to proliferative controls (Fig. S9A), and circadian-range power was significantly increased relative to unreleased cells and UBC-mV-NLS controls (Fig. 5E–F). Upon release, ultradian power decreased (Fig. S9B), dip amplitude increased (Fig. S9C), and mean HES1 protein levels declined, while UBC-mV-NLS levels were unchanged (Fig. 5G).

**Figure 5.**
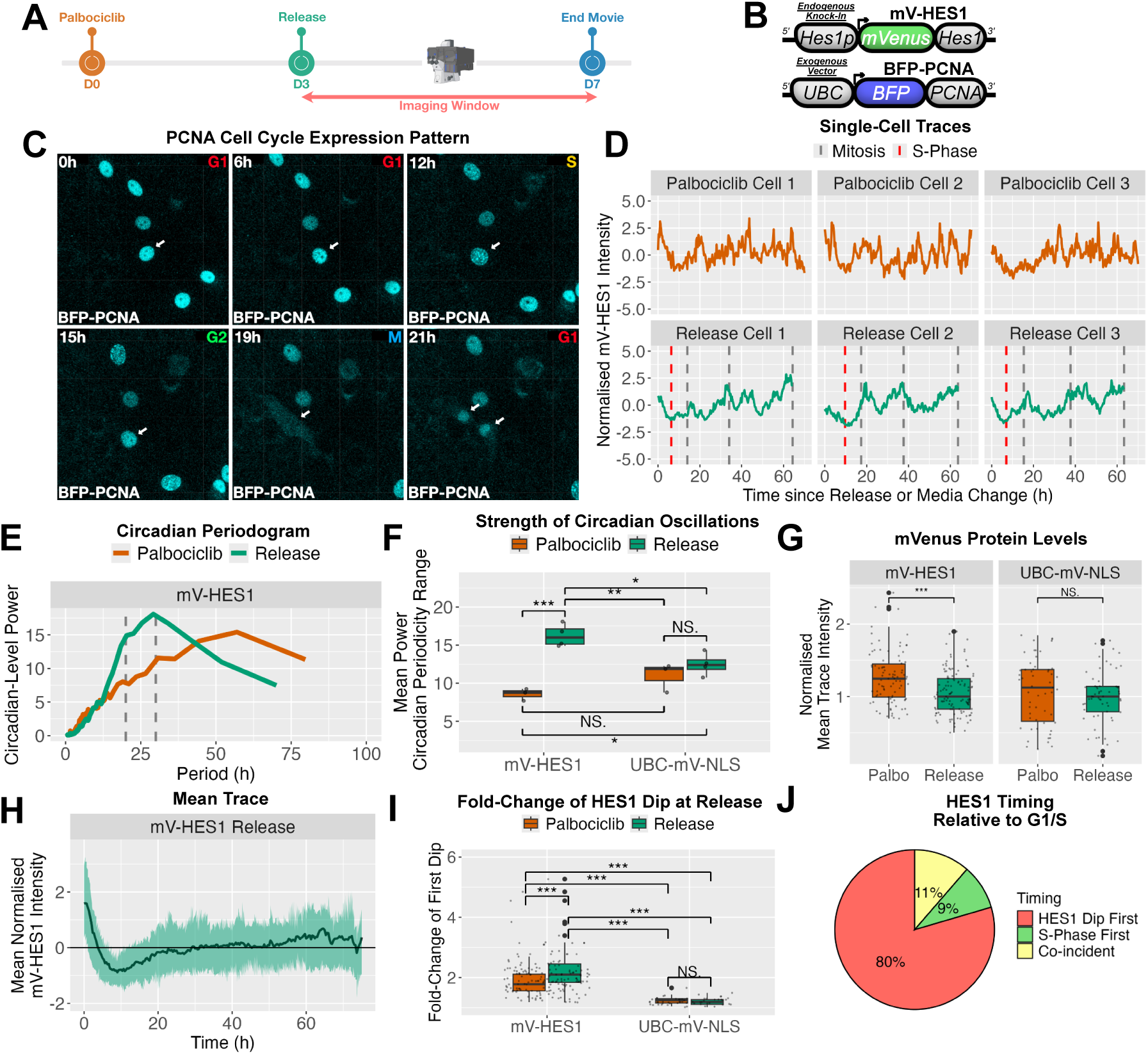
HES1 circadian-level oscillations resume and the G1/S dip is observed upon cell cycle re-entry. (A) Experimental timeline. Cells were arrested with palbociclib for 3 days, released into proliferative media, and monitored by continuous live imaging for a further 4 days. (B) Schematic of constructs used. A UBC-BFP-PCNA reporter was introduced into mV-HES1 cells to mark S-phase via punctate nuclear pattern. (C) Representative snapshots from single-cell live imaging of mV-HES1 UBC-BFP-PCNA cells following release. Arrow highlights a tracked cell progressing through the cell cycle; S-phase is indicated by punctate PCNA at 12h. (D) Representative Z-score–normalised mV-HES1 single-cell time-series from released or continuously arrested cells. For released cells, time is relative to palbociclib withdrawal; for arrested cells, time is relative to a matched media change. Red lines indicate S-phase in the first cell cycle after release. (E) Mean LSP power spectra of circadian-detrended traces from (D). Grey lines indicate the circadian frequency range analysed in (F). N = 3 experiments; n = 115 (Release), 90 (Palbociclib). (F) Mean LSP power within the circadian range (20–30 h) for released versus arrested cells. Each point represents mean power at a given frequency across all cells and replicates. n = 55– 115 cells per group. ANOVA with Tukey’s post-hoc test: ***p < 0.001, **p < 0.01, *p < 0.05. (G) Mean mVenus fluorescence intensity per cell calculated from raw time-series and normalised to the median value from released cells. *N* = 3 experiments, *n* = 55-115 cells per condition. Independent t-test *** < 0.001. (H) Mean Z-score normalised mV-HES1 trace from released cells illustrating the prominent dip following release. Line indicates the mean of *n* = 115 cells pooled from *N* = 3 experiments; shading indicates SD. (I) Fold-change was calculated between mVenus intensity at release and the minimum intensity prior to the 1^st^ mitosis (release cells) or within the first 15h (palbociclib cells). *N* = 3 experiments; *n* = 55–115 cells per group. Independent t-test: ***p < 0.001. Greater fold-change seen in palbociclib mV-HES1 cells than UBC-mV-NLS due to transient response of HES1 to serum during control media change (see SI Methods). (J) Timing of the HES1 dip relative to the G1/S transition based on PCNA signal from (C, D). Events were classified as coincident if occurring within 1h.

Analysis of publicly available RNA-seq datasets [51–52] confirmed reversible regulation of HES1 expression, showing increased HES1 mRNA upon palbociclib treatment (approx. 2-fold) and reduced expression following drug discontinuation in MCF-7 cells (Fig. S10A–B). Thus, these results demonstrate that circadian-level HES1 oscillations are restored upon cell cycle re-entry from arrest, alongside all other parameters that were changed during arrest (ultradian periodicity, dip size and mean levels).

Interestingly, a prominent and reproducible HES1 dip was observed shortly after release (∼8– 10 h post-release) (Fig. 5H), detectable at the population level due to the synchrony of cell cycle re-entry. This initial dip exhibited a mean fold-change of 2.2, significantly greater than dips observed in unreleased cells or UBC-mV-NLS controls during the same interval (Fig. 5I). Co-imaging with the PCNA reporter showed that this dip preceded or coincided with S-phase entry in 89% of cells (Fig. 5J), consistent with the HES1 dip marking the G1/S transition. Thus, cell cycle re-entry is accompanied by a reproducible HES1 dip that can be directly tested for functional significance.

These experiments reveal that circadian-level HES1 oscillations are a hallmark feature of cell cycle progression, which are not observed in arrested cells but can be rescued by re-entry. Moreover, these data establish that the characteristic G1/S HES1 dip is observed during the process of cell cycle re-entry. Taken together, these findings suggest a reciprocal interaction of HES1 and the cell cycle, whereby the cell cycle shapes HES1 dynamics and the HES1 dip observed in proliferative cells may be required for G1/S transition, which was tested next.

### Generation and characterisation of a system for inducible HES1 mis-expression

To examine whether HES1 oscillations (specifically the G1/S dip) are functionally required during cell cycle re-entry, we generated a Tet-On based inducible system to sustain HES1 expression (Fig. 6A). In this system, mScarlet-HES1 is driven by a Tet-responsive promoter (TRE), herein called the Tet-mS-HES1 cassette, which can be precisely induced by doxycycline treatment. This exogenous cassette was expressed in endogenous mV-HES1 MCF-7 cells, such that expression of each HES1 reporter can be distinguished by the fluorophore. A Tet-driven mScarlet-NLS reporter (Tet-ms-NLS) served as a control.

**Figure 6.**
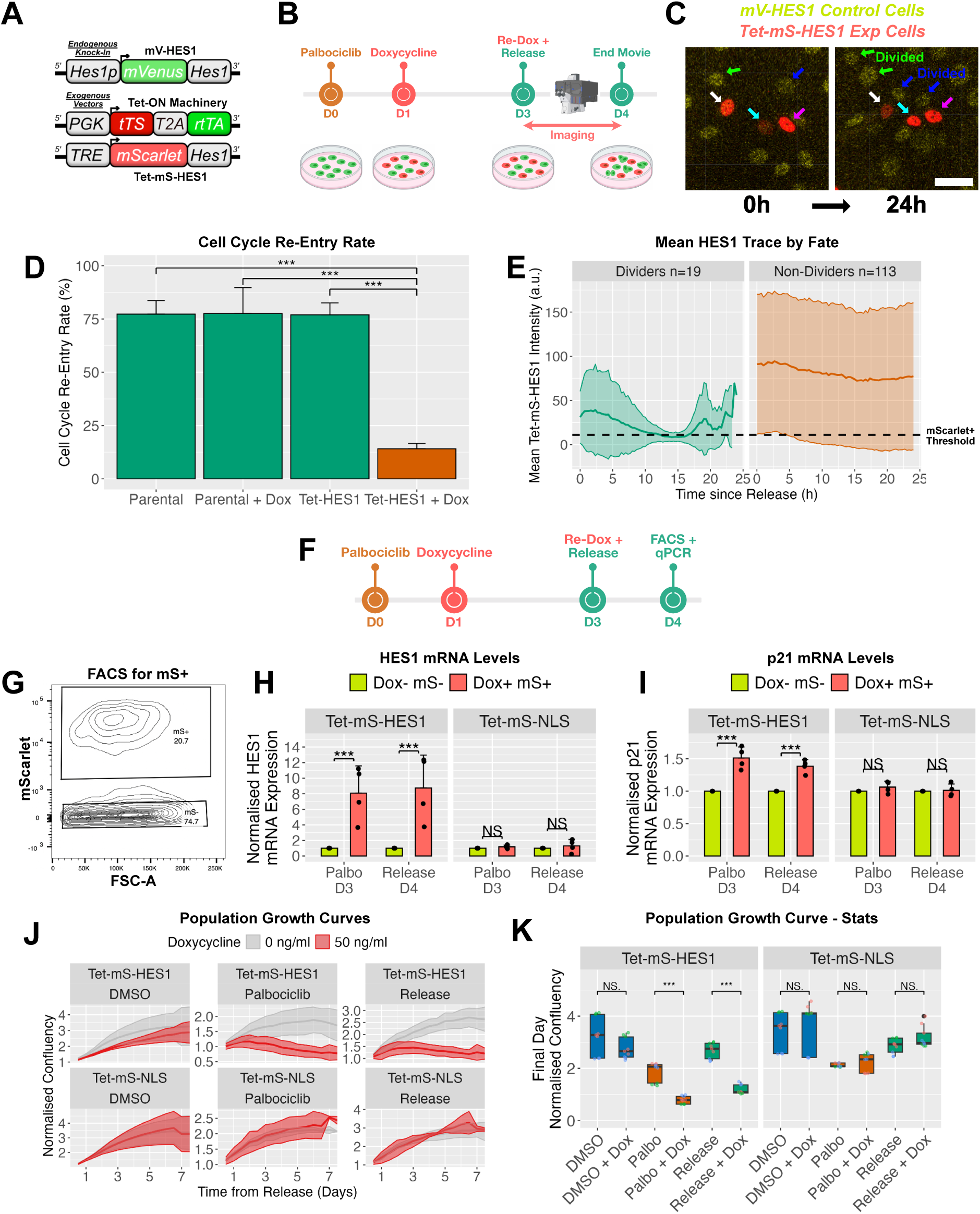
Induced expression of less-dynamic HES1 during release upregulates p21, impedes cell cycle re-entry and prevents population outgrowth. (A) Schematic of constructs used. Tet-On lentiviral vectors (see methods) were incorporated into mV-HES1 cells to enable doxycycline-inducible Tet-mS-HES1 expression. (B) Timeline of the 24h release assay used in (C-H). Cells were arrested with palbociclib for 3 days, then released into proliferative media ± doxycycline, and imaged continuously for 24 h to assess cell cycle re-entry. Cartoon illustrates a typical field: red and green nuclei indicate cells that did or did not induce Tet-mS-HES1, respectively. (C) Representative time-lapse images from the 24h release assay showing cells at 0h (release) and 24h. Arrow colours mark distinct cell lineages. Control cells without Tet-mS-HES1 expression typically divided (green and blue arrows), whereas Tet-mS-HES1–positive cells did not (white, cyan, magenta arrows). Scale bar = 40 μm. (D) Quantification of the proportion of cells undergoing mitosis during the 24 h release assay. Parental and Tet-HES1 refer to mV-HES1 cells without or with the Tet-mS-HES1 cassette, respectively. Doxycycline = 5 ng/mL. Cells that died during imaging were excluded. Bars show mean ± SD from N = 3 experiments; n = 130–180 cells per group. Independent t-test: ***p < 0.001. (E) Single-cell Tet-mS-HES1 time-series from doxycycline-treated cells in (D) were grouped by division fate (Dividers vs Non-dividers) and averaged into a mean trace. The threshold for Tet- mS-HES1 positivity is indicated. (F) Experimental timeline for (G–I). Cells expressing Tet-mS-HES1 or control Tet-mS-NLS were arrested ± doxycycline and sorted by FACS to isolate mScarlet⁺ cells either during arrest (day 3) or 24 h post-release (day 4). (G) Representative FACS plot displaying sorting of mScarlet+ (mS+) cells. (H, I) Sorted mScarlet+ cells were analysed by RT-qPCR for expression of (H) HES1 and (I) p21 transcripts. Bars show mean ± SD from *N* = 4 experiments. Values are fold-change from the Dox- mS- sample. ANOVA with Tukey’s post-hoc test. ***p < 0.001, **p < 0.01. (J) Seven-day population growth curves following release. Cells expressing Tet-mS-HES1 or Tet- mS-NLS were monitored under proliferative (DMSO), arrested (palbociclib), or released conditions ± doxycycline (50 ng/mL). Lines show mean normalised confluency; shading indicates SD. N = 3 experiments. (K) Quantification of growth curves from (I). Confluency measurements from the final day (3 time-points per day per experiment) were pooled. ANOVA with Tukey’s post-hoc test: ***p < 0.001.

Because HES1 autorepresses its own promoter, we anticipated that Tet-driven exogenous HES1 would remain constant and thereby impose continuous repression on endogenous HES1 [38]. Instead, Tet-mS-HES1 itself unexpectedly oscillated at the single cell level (Fig. S11A), most likely due to a combination of promoter dynamics and protein instability (see also [53]), creating periodic windows in which endogenous HES1 escaped repression. This gave rise to pronounced anti-phase oscillations between the endogenous and exogenous reporters in single cells.

We reasoned that such anti-phase behaviour could reduce the time that total HES1 spends in a low-expression state, because when the exogenous expression is low, the endogenous HES1 is expressed, effectively producing a less dynamic overall profile. To test this, we performed immunostaining for total HES1 and compared the coefficient of variation between wild-type cells (endogenous HES1 only) and doxycycline-induced Tet-mS-HES1 cells (expressing endogenous and exogenous HES1 in anti-phase) (Fig. S11B). Indeed, Tet-mS-HES1–positive cells exhibited reduced population-level variance in total HES1 compared to controls, consistent with a less dynamic, more sustained total HES1 state, likely arising from anti- correlated dynamics between endogenous and exogenous HES1 in single cells (Fig. S11C).

### Mis-expression of HES1 during cell cycle release impedes cell cycle re-entry

To investigate whether the HES1 dip is necessary for cell cycle re-entry, Tet-mS-HES1 expression was induced at the point of release from palbociclib (Fig. 6B), thereby maintaining HES1 levels specifically as cells attempt to re-enter the cell cycle and HES1 would ordinarily dip. Moreover, anti-correlated expression of Tet-mS-HES1 and endogenous mV-HES1 was observed in time-series from this assay, consistent with less dynamic cumulative expression (Fig. S11D). This markedly impaired cell cycle re-entry, reducing the fraction of cells dividing within 24h of release from 75% in controls to 14% in Tet-mS-HES1–positive cells (Fig. 6C–D).

Among the small fraction of Tet-mS-HES1–expressing cells that did divide, a synchronous decline in HES1 was observed, producing a visible dip in the population mean at ∼12–15h post-release (Fig. 6E), reminiscent of the endogenous G1/S-associated HES1 dip. In contrast, cells that failed to re-enter exhibited a largely flat mean HES1 profile. This shows that a consistent dip in total HES1 is observed when a cell successfully re-enters the cell cycle but not in cells which are impeded, supporting a requirement for this dip in efficient re-entry.

To explore how HES1 mis-expression impedes re-entry, we examined the G1/S regulator and cell cycle inhibitor p21, a known HES1 target that has been shown to be upregulated under sustained HES1 expression [37–38]. Cells were arrested with palbociclib and treated with doxycycline to induce Tet-mS-HES1 (Fig. 6F), after which mScarlet-positive cells were isolated by FACS either during arrest or following release for qPCR analysis (Fig. 6F–G). HES1 transcripts were strongly upregulated (8–9 fold) in Tet-mS-HES1 cells, but not in Tet-mS-NLS controls (Fig. 6H). p21 expression was also reproducibly elevated in Tet-mS-HES1 cells during arrest (1.5 fold) and 24h post-release (1.4 fold), but unchanged in controls (Fig. 6I). These data suggest that HES1 mis-expression promotes p21 upregulation, which may contribute to impaired cell cycle re-entry following palbociclib withdrawal.

### HES1 mis-expression suppress long-term growth and promote cell death

We next asked whether reduced HES1 dynamics exert long-lasting effects beyond the initial 24h re-entry window. To test this, we performed population-level growth curve analysis for 7 days following palbociclib withdrawal (Fig. 6J–K). Induction of less dynamic HES1 had no significant effect on the growth of proliferative (DMSO-treated) cells. In contrast, inducing less dynamic HES1 at the time of release from palbociclib prevented population outgrowth and produced negative growth slopes from approximately day 3 onward (Tet-mS-HES1, release; Fig. 6J). A similar negative growth trajectory was observed when less dynamic HES1 was induced during continued palbociclib treatment (Tet-mS-HES1, palbociclib; Fig. 6J). These negative growth rates are suggestive of a net reduction in cell number due to cell death.

Consistent with this interpretation, we observed a dose-dependent increase in morphologically-defined cell death in released Tet-mS-HES1 cells exposed to escalating doxycycline concentrations (Fig. S12A–C). Molecular viability assays further revealed a reduction in viable cells from 80–83% in controls to 52% upon Tet-mS-HES1 induction, driven by increases in both apoptotic and necrotic populations (Fig. S12D–G). By contrast, doxycycline treatment of control Tet-mS-NLS cells had no effect on population growth (Fig. 6J– K) or cell viability (Fig. S12D–G). Together, these results suggest that that perturbation of HES1 dynamics at the time of release from G0/G1 arrest impairs cell cycle re-entry, induces cell death and thus limits subsequent population expansion.

We next sought to investigate the role of HES1 in breast cancer patients, using publicly available datasets [54]. We found, high levels of HES1 are significantly associated with longer relapse-free survival of both luminal A and ER+ breast cancer patients (Fig. S13A-B). This prolonged survival is consistent with our findings that HES1 overexpression impedes cell cycle re-entry and induces cell death in arrested cells. This may point to higher, or sustained, levels of HES1 preventing re-activation from dormancy and subsequent relapse in ER+ breast cancer patients.

## Discussion

The work described here addresses the clinical problem of tumour recurrence in ER⁺ breast cancer by focusing on a fundamental, cell-autonomous process: re-entry of quiescent cancer cells into the cell cycle. While recurrence is a multifactorial phenomenon involving interactions between disseminated tumour cells (DTCs) and the tumour microenvironment, cell cycle reactivation of dormant cells is a necessary initiating step. Although reactivation can occur stochastically [8], we specifically modelled re-entry following withdrawal of the CDK4/6 inhibitor palbociclib. This is clinically relevant because CDK4/6 inhibitors are widely used to suppress proliferation in ER⁺ breast cancer, yet tumours ultimately escape their cytostatic effects. While our system involves drug withdrawal, the mechanisms identified here are likely more broadly relevant to emergence from quiescence.

Extending previous work [39], we show that the Notch target HES1 exhibits nested oscillatory dynamics in proliferative MCF-7 cells, comprising a dominant ∼24 h oscillation and a faster ultradian component (∼6–7 h). The ∼24 h oscillation matches cell cycle duration, is present in proliferative cells, lost during arrest, and restored upon re-entry, indicating that it is paced by cell cycle progression. In contrast, ultradian oscillations show no reproducible association with cell cycle phase and are not detectably distinct from noise in proliferative cells. We therefore propose that these faster oscillations arise autonomously from HES1 autorepression combined with mRNA and protein instability, a mechanism known to generate ultradian oscillations in other systems [55–59]. Their increased prominence during arrest is likely a secondary consequence of the loss of the dominant cell-cycle–linked oscillation, consistent with general properties of nested oscillatory systems in which attenuation of the slower dominant rhythm enhances the prominence of faster autonomous components [60].

In developmental systems, HES1 oscillations can be cell autonomous [57], but in a multicellular environment they are likely coupled between cells via Notch signalling, which provides a positive input to HES transcription [61,62]. Notch signalling can link HES1 regulation not only to environmental cues but also to the cell cycle. Notably, the Notch intracellular domain (NICD) is phosphorylated by CDK2 and degraded during S phase [63], which could transiently interrupt Notch-driven HES1 transcription and generate the characteristic G1/S dip in HES1 protein levels. In addition, direct post-translational regulation of HES1 by the cell cycle machinery has been described in Xenopus, where phosphorylation by CDK4 and CDK2 promotes HES1 degradation [64]. Together, these mechanisms provide a plausible explanation for the inverse relationship we observe between CDK2 activity and HES1 levels, as well as for the loss of circadian-level oscillations and elevation of HES1 protein levels following palbociclib treatment, which inhibits both CDK4/6 directly and CDK2 indirectly [65].

Beyond demonstrating that circadian-level HES1 oscillations are driven by the cell cycle, our data indicate that these oscillations also feedback to regulate cell cycle progression, as predicted by theoretical models [66]. This is supported by our finding that preventing the HES1 dip normally observed prior to S-phase entry impairs cell cycle re-entry from arrest.

A key mechanistic question arising from this work is how distinct HES1 dynamics differentially regulates the cell cycle. HES1 transcriptionally represses both cell cycle activators (CycD1, CycE2, CycA2 and E2F) and inhibitors (p21, p27) [25–29,67]. In proliferating cells, oscillatory HES1 expression may permit periodic de-repression of cyclins during the dip preceding S-phase, while elevated HES1 levels in G1 may suppress p21 and p27 to facilitate progression through G1/S [47,48].

By contrast, sustained HES1 may continuously repress cyclins, thus preventing cell cycle progression. Indeed, sustained HES1 has been shown in murine neural stem cells to indirectly upregulate p21 via repression of a p21 repressor [37,38], and we similarly observe p21 induction during HES1 mis-expression here. Thus, we surmise sustained HES1 may both repress cyclins and activate p21. Together, these effects would impose a dual blockade on cell cycle re-entry, maintaining arrest even after CDK4/6 inhibitor withdrawal, as occurs clinically during resistance. Consistent with this model, spontaneously quiescent cells exhibited HES1 dynamic changes similar to those induced by palbociclib, suggesting similar vulnerabilities may be common across quiescence.

Because the Tet-On HES1 system alters both the temporal dynamics and the overall levels of HES1, it is difficult to fully disentangle the relative contributions of each perturbation. Notably, however, during both palbociclib-mediated and spontaneous arrest, mean HES1 protein levels increase only modestly (∼1.4-fold) and remain within the physiological range observed in proliferative cells, indicating that HES1 levels do not have to be dramatically increased during arrest.

In addition to impeding cell cycle re-entry, mis-expression of HES1 induced cell death. Whether this reflects a direct effect of HES1 on cell death pathways or arises indirectly from cellular stress associated with HES1-mediated prolonged arrest remains unclear. However, increased stability of the HES1 orthologue has been shown to induce apoptosis via a p53- dependent mechanism in Xenopus neural development [68], raising the possibility of a conserved pathway. Given that p53 itself exhibits oscillatory dynamics in MCF-7 cells [69], future work examining the interplay between HES1 and p53 dynamics may be informative.

Importantly, less dynamic HES1 expression induced negative population growth rates in both arrested cultures and following release from arrest. This may be of significant clinical benefit, as current therapies primarily target proliferative cancer cells. Manipulating HES1 dynamics could therefore represent a strategy to eliminate quiescent tumour cells, reduce residual disease, and limit reactivation. Nonetheless, as these findings were obtained in simplified 2D in vitro systems, validation in more physiologically relevant models, such as patient-derived xenografts, will be essential before translational application.

In conclusion, we found that the expression dynamics of HES1 changed dramatically and reversibly during entry into and exit from cell cycle arrest. These changes are functionally important, as their perturbation prevented reactivation. Thus, our findings suggest that targeting protein dynamics may offer a promising and precise strategy to prevent emergence from dormancy in breast cancer.

## Materials and Methods

Briefly, ER⁺ (MCF-7) and triple-negative (SUM149) breast cancer cell lines were engineered with fluorescent fusion reporters for HES1 and other cell cycle regulators using CRISPR–Cas9– mediated knock-in and lentiviral integration. Cell cycle arrest and re-entry were modelled using reversible CDK4/6 inhibition with palbociclib. Single-cell protein dynamics were quantified by long-term live-cell imaging using confocal microscopy, in combination with fluorescent reporters of HES1, CDK2 activity, p21 expression and PCNA. Time-series were analysed using established signal processing approaches, including Lomb–Scargle periodograms and autocorrelation analysis, to characterise oscillatory behaviour.

To perturb HES1 dynamics during cell cycle re-entry, a doxycycline-inducible Tet-On system was used to drive exogenous HES1 expression. The effects on proliferation and cell cycle re-entry, were assessed by live-imaging, immunofluorescence microscopy, flow cytometry, and RT-qPCR. Cell cycle arrest, re-entry, and cell death were quantified using population growth assays, mitotic tracking, PCNA-based reporters, Ki67 index, EdU incorporation and viability analyses. Bioinformatic analyses of publicly available transcriptomic datasets were used to validate key observations. Full experimental procedures, imaging parameters, data analysis pipelines, statistical methods, and reagent details are provided in the SI Appendix, Materials and Methods.

## Acknowledgements

The authors would like to thank Dave Spiller, James Bagnall and Miruna Verdes from the Bioimaging Facility, FBMH, University of Manchester; David Chapman, Bradley Dean and Gareth Howell from the Flow Cytometry Facility, FBMH; All members of the Papalopulu and Clarke labs for their comments and discussion. The work was supported by a Cancer Research UK—Engineering and Physical Research Sciences Research Council Multidisciplinary Award grant to N.P. and R.B.C. (DRCMDP-Nov22/100009), a Wellcome Trust Investigator Award to N.P. (224394/Z/21/Z) and a Medical Research Council PhD studentship to O.C.

