## Supplementary material for "HES1 oscillations are required for cell cycle re-entry in oestrogen receptor positive breast cancer cells": SI Appendix

### Supplementary Materials and Methods

#### Cell Culture

MCF-7 cells were cultured in high-glucose DMEM (D6429, Sigma) supplemented with 10% heat-inactivated FBS (10500064, Gibco). SUM149 cells were cultured in Ham's F-12 (N6658, Sigma), supplemented with 5% FBS, 5 ug/mL insulin (I6634, Sigma), and 1 ug/mL hydrocortisone (H0888, Sigma). All cells were maintained at 37°C with 5% $CO_2$ in a humidified incubator. Cells were routinely tested for mycoplasma and typically passaged twice per week.

To induce cell cycle arrest, MCF-7 cells were treated with 1 µM palbociclib (PZ0383, Sigma) or an equal volume of DMSO (D8418, Sigma) for 3–4 days. SUM149 cells were treated with 10 µM palbociclib. For release, cells were washed with pre-warmed PBS (2–3 min) and replaced into growth media containing DMSO instead of palbociclib. Doxycycline was added at 5 or 50 ng/mL (as indicated) (D9891, Sigma), 24 h after palbociclib addition (Day 1) and again at release (Day 3). Unreleased controls received fresh palbociclib-containing media.

The addition of fresh palbociclib-containing media occasionally produced minor artefactual effects, by virtue of providing fresh serum, which induced a transient proliferative stimulus despite the addition of extra palbociclib. This resulted in higher proliferative index in methodologies which included a media change in the palbociclib condition (Fig. 1D) compared to those which did not (Fig. 1C). Moreover, HES1 is a serum-induced gene (Yoshiura *et al*., 2007) so unreleased controls receiving fresh palbociclib medium exhibited a transient peak in HES1 levels due to the fresh serum, manifesting in a subsequent dip-phase, which was nonetheless smaller than the dip in released cells (Fig. 5I), which additionally arises from HES1 dynamics at G1/S.

#### Reporter Constructs and Cell Lines

The constitutive nuclear mVenus reporter UBC-mV-NLS, driven by the Ubiquitin C promoter, was used as a live-imaging control (Sabherwal et al., 2021). The PCNA-based S-phase reporter from our previous study (Sabherwal et al., 2021) was modified by replacing mCherry with mTagBFP2 to generate UBC-BFP-PCNA. A reporter of CDK2 activity comprising of CMV driving the CDK2 sensor DHB-mScarlet and the nuclear marker H2B-miRFP670, separated by P2A (DHB-mScarlet-P2A-H2B-miRFP670) was modified from an existing construct (Adikes *et al*., 2020) to introduce a mammalian promoter and optimise fluorescent channels. A reporter of HES1 dynamics that could be exogenously introduced into various cell lines was commercially acquired (VectorBuilder), comprising the 2.9kb human HES1 promoter driving mScarlet fused to HES1 cDNA and 3' UTR (HES1p-mS-HES1).

To modulate HES1 expression dynamics, we used a second-generation Tet-On system (VectorBuilder), comprising two lentiviral plasmids:

1. PGK-tTS-T2A-rtTA — expressing the tetracycline transcriptional silencer (tTS) and the reverse tetracycline-controlled transactivator (rtTA), separated by a T2A peptide under the PGK promoter. Also contains blasticidin resistance gene driven by PGK.
2. TRE-mS-HES1 or TRE-mS-NLS — expressing mScarlet (mS) fused to either full-length HES1 or a nuclear localization signal, under control of the tetracycline-responsive element promoter (TRE). Also contains neomycin resistance gene driven by PGK.

MCF-7 cells previously engineered via CRISPR-Cas9 to express an mVenus-HES1 fusion protein (mV-HES1) (Sabherwal et al., 2021) were further modified to express UBC-BFP-PCNA, PGK-tTS-T2A-rtTA, and either TRE-mS-HES1 or TRE-mS-NLS, yielding Tet-mS-HES1 and Tet-mS-NLS cell lines. Parental MCF-7 cells were also transduced with UBC-mV-NLS to generate a control line. Additionally, DHB-mScarlet-P2A-H2B-miRFP670 was transiently incorporated into mV-HES1 cells to visualise relative dynamics of HES1 and CDK2 actiivty. MCF-7 cells expressing an endogenous p21-mVenus fusion-reporter (Stewart-Ornstein *et al*., 2016) were further engineered to express the exogenous HES1p-mS-HES1 reporter, by lentiviral incorporation, so that p21 and HES1 could be simultaneously analysed.

#### Generation of mScarlet-HES1 Endogenous Fusion Reporter in SUM149 cells

HES1 in SUM149 cells was endogenously tagged at the N-terminus with in-frame mScarlet-I using CRISPR/Cas9 genome-editing technology (mS-HES1). The donor plasmid for endogenous tagging was commercially synthesised (GenScript), featuring a cassette consisting of ATG-3XFLAG mScarlet-linker, flanked by 800 bp homology arms upstream (5' homology arm) and downstream (3' homology arm) of the HES1 start codon. The plasmid resides in a pUC57 based vector backbone. gRNAs against human Hes1 were designed using the Wellcome Trust Sanger Institute portal (https://wge.stemcell.sanger.ac.uk//). gRNAs, tRNA, and purified Sp-Cas9 proteins were obtained commercially (Integrated DNA Technologies). The gRNA 5'-GAAAAATTCCTCGTCCCCGGTGG-3' was used for directing the CRISPR-Cas9 reaction. The mScarlet cassette was inserted in-frame, downstream of the endogenous HES1 ATG through homology-directed repair (HDR), using donor plasmid as a template. Shield mutations were introduced in the donor plasmid, near the PAM site, to abrogate repeat cleaving of the edited template (Fig. S7).

A gRNA (48μM) and tRNA (48μM) mixture was heated at 95°C for 5 min, followed by slow cooling at room temperature. To make the RNP complex, 5μL of gRNA:tRNA complex was mixed with 2μL of Cas9 (61μM) and 4μL of IDT duplex buffer, resulting in final concentrations of both gRNA and tRNA as 20.4μM, and Cas9 as 11.09μM. The RNP complex was incubated at room temperature for 10 min prior to being mixed with 2μg of donor plasmid. The RNP-donor plasmid mix was used to electroporate one million cells, resuspended in 100 μL of Lonza P3 Primary Cell Nucleofector buffer (Lonza). The EN-130 program on the 4D-Nucleofector X Unit (Lonza) was used for electroporation. Post electroporation, cells were cultured in growth media containing 1 μM Alt-R™ HDR Enhancer (1007910, IDT). HDR Enhancer was removed from media after 24 hours. Electroporated cells were FACS-sorted against mScarlet fluorescence, and single-cell clones were grown in Ham's F-12 media, on wells coated with vitronectin at 0.5ug/cm$^2$. Clones were expanded and genotyped to identify successful knock-ins. A total of 10 lines were genotyped and a homozygous clonal cell line (P1C10) was then taken forward. TOPO-cloning was used to sequence both alleles of P1C10 to confirm there were no undesired mutations (Primers: FP1 for 5' and RP1 for 3'). Primers used for genotyping and sequencing are detailed in supplementary table S1.

**Lentivirus Production and Transduction**

Reporter constructs were packaged into lentivirus using HEK-293-LTV cells (LTV-100, Biolabs). Two T75 flasks were seeded with 700,000 cells each and cultured for 72 h. Transfections were performed by combining 54 μL 1× PEI (919012, Merck) in 500 μL serum-free media (D1145, Sigma) with a DNA mix containing 6 μg MD2.G, 9 μg psPax2, and 12 μg lentiviral expression plasmid in another 500 μL of serum-free media. After 2 min (PEI) and 30 min (combined mix) of incubation at room temperature, 500 μL of the final solution was added dropwise to each flask. After 24h, media was replaced with 10 mL fresh growth medium supplemented with 10 mM sodium butyrate (19137, Sigma) and incubated for 6–8h. Medium was then replaced again, and virus-containing supernatant was collected 40–48h later, filtered (0.45 μm), and mixed with 5 mL 5× PEG virus precipitation solution (LV810A-1, SBI). The solution was incubated at 4°C for 24–96 h, centrifuged at >1,500 × g for 30–60 min, and resuspended in 200 μL chilled PBS. Aliquots were stored at –80°C.

For transduction, 100,000 MCF-7 cells were seeded per well in a 6-well plate, then infected with one aliquot of virus in 1 mL media containing 0.1% polybrene (TR-1003-G, Sigma). After 24–48h, cells were selected with 20 μg/mL blasticidin (15205, Sigma) or 5 mg/mL G418 (4727878001, Sigma) for 72h, followed by recovery in standard growth media. Fluorescent populations were enriched by FACS using a Bigfoot Spectral Cell Sorter (Thermo) operated by the Flow Cytometry Core Facility, University of Manchester.

**RNA Extraction and RT-qPCR**

Total RNA was generally extracted using the RNeasy kit (74104, Qiagen). For FACS-isolated samples sorted on mScarlet expression, where cell numbers were limited, phenol-choloroform based extraction was performed to optimise yield, as previously described (Miller *et al*., 2025). Next, 250 ng of RNA was reverse-transcribed using SuperScript III Reverse Transcriptase (18080044, Thermo). Resulting cDNA was diluted 1:10, corresponding to ~25 ng RNA per qPCR reaction. qPCR was performed using Power SYBR Master Mix (4367659, Thermo). Primer pairs are listed in supplementary table S1. Only technical replicates within 0.1 CT of each other were included in the analysis. Gene expression was normalized to housekeeping genes (GAPDH and ActinB/Rpl7) by calculating ΔCT, and relative expression was determined as 2^–ΔCT. Where indicated, values were further normalized to the median of a designated control condition to estimate fold change.

**Immunostaining and Immunofluorescence Microscopy**

Cells were seeded on glass coverslips and treated with drugs as indicated. At endpoint, cells were fixed with 4% PFA/PBS and permeabilised using 0.5% Triton X-100 in PBS (+ 10% donkey serum, D9663 Sigma, when donkey antibodies were used). Primary antibody incubation was performed overnight at 4°C, followed by secondary antibody incubation for 2h at room temperature in the dark. Antibodies were diluted in 1× Western Blocking Reagent (Sigma) in PBS. Nuclei were stained with 1 µg/mL DAPI (62248, Thermo), and coverslips were mounted using ProLong Diamond Antifade Mountant (P36934, Invitrogen). Primary antibodies for mouse anti-Human Ki67 (1:100, 550609, BD Biosciences), mouse anti-Human HES1 (1:50, SC-166410, Clone E5, Santa Cruz), rat anti-RFP (1:500, 5F8-150, Chromotek) were used. Secondary antibodies were goat anti-mouse Alexa Fluor 568 (1:1000, A11004, Invitrogen), donkey anti-mouse Alexa Fluor 647 (1:1000, A31571, Invitrogen) and donkey anti-rat DyLight 550 (1:800, SA5-10027, Invitrogen).

Images were acquired using an Olympus IX83 inverted microscope (10× objective) equipped with an Orca ER camera (Hamamatsu) and CellSens software (Olympus). Custom Fiji macros were used for batch image preprocessing. Quantification of Ki67+ nuclei was performed using a MATLAB pipeline (Marinopoulou et al., 2021) involving DAPI-based segmentation, followed by intensity extraction per nucleus. Ki67+ cells were defined as those exceeding the fluorescence threshold set by secondary-only controls.

For Coefficient of Variation (CV) analysis, images were segmented on DAPI using Cellpose-SAM (Pachitariu et al., 2025) and the resulting masks were then used to extract values for total nuclear HES1 and mScarlet intensity. This was done using the regionprops-table function in the scikit-image python library (van der Walt et al., 2014). mScarlet+ cells were thresholded based on the maximum expression value in the control sample (excluding any outliers). CV was calculated by dividing the standard deviation of total HES1 signal in each sample by the mean value. CV was then compared between doxycycline-treated mScarlet+ cells and mScarlet- controls.

**IncuCyte Imaging and Population Growth Analysis**

Cells were seeded at 2,850 cells/cm² in 12-well plates and treated with DMSO, palbociclib, or doxycycline (D9891, Sigma) as required. Immediately following treatment, plates were placed into the IncuCyte Zoom live-cell imaging system (Essen Biosciences). Whole-well 4× phase contrast images were acquired every 3–12h over 10–14 days. Treatments and releases were performed in tissue culture between scans. All wells were subject to the same handling, including media replacement and washes, to control for serum-related effects.

Image analysis was conducted using IncuCyte software, where custom cell line-specific segmentation masks were applied to estimate percentage confluency. Confluency was normalised either to the initial seeding density or to the confluency at release to allow fold-change comparison. Growth curves represent the mean of experimental replicates, with shaded areas indicating standard deviation. To test statistical significance, confluency measurements from the final 24h of the time course were pooled and compared across conditions.

**EdU Incorporation Assay**

MCF-7 cells were seeded at 5,200 cells/cm² in 6-well plates and treated with either DMSO, palbociclib continuously, or palbociclib for 3 days followed by release. On Day 3, 2.5 µM EdU (5-Ethynyl-2'-deoxyuridine) was added during the media change or release. After 24h, cells were harvested and processed for EdU detection using the Click-iT Plus Flow Cytometry Kit (C10634, Thermo), following the manufacturer's instructions. Samples were analysed using an LSRFortessa SORP flow cytometer (BD Biosciences).

**Cell Death Assay**

To assay viability, 500,000 cells were harvested and stained with Zombie NIR (423105, Biolegend) at a dilution of 1:5000, according to manufacturer's instructions. Next, cells were washed with Cell Staining Buffer (420201, Biolegend), stained with APC-Annexin V (640920, Biolegend) and washed with Annexin V Binding Buffer (422201, Biolegend). Stained samples were analysed analysed using an LSRFortessa SORP flow cytometer (BD Biosciences). Cells were then designated as viable (Zombie- Annexin-), necrotic (Zombie+ Annexin-) or apoptotic (Zombie+/- Annexin+).

**Quantitative Single-Cell Live Imaging and Tracking**

Cells were seeded in 4-compartment glass-bottom dishes (627870, Greiner) and imaged within the chamber of a Zeiss LSM880 inverted confocal microscope maintained at 37 °C and 5% $CO_2$. Using a 20× objective with 0.6x zoom, 2×2 grid live images were acquired every 20 min for up to 100h, depending on the experiment. Z-stacks spanning ~40 µm (typical nuclear signal range: 30–35 µm) were captured and collapsed via maximum intensity projection. Resulting time-lapse movies were manually tracked using Imaris (Bitplane) by placing 9 µm diameter spots (to cover majority of nucleus) over individual cells to capture nuclear fluorescence. Manual tracking was utilised due to the necessity to track cells over divisions, which is often poorly performed by automated approaches. Moreover, due to oscillations of HES1, nuclear signal intensity often reached a nadir which automated algorithms struggled to detect. Cells were typically tracked from mitosis to mitosis, to identify cell cycle periodicities, unless stated otherwise. For tracking of the CDK2 sensor (DHB-mScarlet), a spot was placed over both the nucleus and the cytoplasm, so that a cytoplasmic:nuclear ratio could be calculated, as higher ratio values are indicative of greater CDK2 activity (Adikes *et al*., 2020).

DMSO-treated control cells were tracked for at least two full generations to capture recurring dynamics. In palbociclib-treated conditions, cells were tracked for the full movie duration, rare dividing cells under palbociclib arrest conditions were excluded from the main cohort and analysed separately. In release experiments, cells were tracked from the first frame until at least two mitoses post-release were observed. Tracking data including x, y, z coordinates, mean nuclear fluorescence intensity, and time were exported from Imaris and processed using *tRecs*, a custom Python script (https://github.com/TMinchington/tRecs), which provides information on lineages and mitotic events. Aggregated master files were then imported into MATLAB or RStudio for downstream analysis.

**Time-Series Visualisation and Protein Level Estimation**

Nuclear protein levels were represented by the mean fluorescence intensity of each manually placed spot. Time-series were imported into RStudio and visualisation was performed using *ggplot2* from the *tidyverse* library (https://www.tidyverse.org/). Individual time-series were predominantly Z-score normalised per cell (mean trace intensity value was subtracted from the raw value at each timepoint and divided by the standard deviation). This normalisation was performed to control for differences in fluorescent intensity levels between individual cells. For comparison of expression dynamics between mVenus and mScarlet reporters in the same cell, each trace was additionally min–max scaled to a 0–1 range based on that cell's minimum and maximum expression values, allowing relative trace shapes to be visualised on the same scale. The time plotted refers to time since the track started rather than when the movie started so is independent to each cell. This is predominantly the time since the first mitosis of the track for proliferative cells, or the start of the movie for palbociclib-arrested and released cells. S-phase events were manually annotated based on punctate UBC-BFP-PCNA signals (Zerjatke *et al*., 2015) and incorporated into trace metadata.

To assess population-wide dynamics, mean traces were constructed by aggregating Z-scored values and plotting the average with standard deviation across aligned timepoints. Additionally, individual Z-scored traces were visualised as heatmaps using *geom_tile* in *ggplot2*, where each column represented a cell and colour denoted fluorescence intensity. For DMSO-treated two-generation tracks, time was aligned to the third and final mitosis; for palbociclib-arrested cells, alignment was to the endpoint of the time series.

Protein abundance between conditions was estimated by calculating the mean raw nuclear fluorescence intensity of each cell trace in RStudio and visualised via box plots across experimental replicates. Minimum and maximum intensities for each trace were also quantified.

**Spontaneous Non-Divider Dataset**

mV-HES1 cells were cultured in standard growth media and continuously live-imaged for up to five days. Amidst these proliferative conditions, a subset of cells exhibited extended periods without division, including some that remained non-proliferative throughout the entire time-course. Cells that failed to divide for at least 60 hours, exceeding the average duration of two generations (57h), were classified as spontaneous non-dividers (SNDs). Time-series from SNDs were compared to mother-daughter traces from actively cycling cells from the same culture and analysed for differences in mV-HES1 periodicity.

**Single-Cell Live Imaging Re-Entry Assay**

To quantify single-cell division events a live-imaging assay was performed. The number of cells present at the start of the time-course was manually counted. Divisions were manually assigned based upon entry into mitosis phase, characterised by translocation of mV-HES1 from the nucleus to the cytoplasm, due to nuclear envelope breakdown (as in Fig. 1E). In cells released from palbociclib, a division was classified as a re-entry event. In the release condition, only the first division per lineage was counted as a re-entry event to avoid false inflation from subsequent divisions within the same lineage. For example, a cell dividing at 1h and again at 23h post-release would otherwise be misclassified as two independent re-entries. To ensure that only genuinely arrested cells were included in the re-entry rate analysis, cells that divided within the first 8h post-release were excluded, because the known 10–14 h G1/S-to-mitosis interval in these cells (Sabherwal *et al.*, 2021) indicates that cells dividing this early must have already progressed beyond G1/S prior to release and were therefore incompletely arrested.

In the Tet-mS-HES1 re-entry assay, cells were classified as Tet-mS-HES1 positive if their mScarlet fluorescence exceeded a threshold set by the maximum extra-cellular background intensity in the image. All mScarlet-positive were then followed for 24h (based on the average time of re-entry defined in the earlier assay) and manually assigned to one of three potential fates: division, no division or death. Cell death was identified based on characteristic morphology and behaviour, including increased autofluorescence, loss of motility, nuclear shrinkage, membrane blebbing, and cessation of proliferation. In control cell lines lacking Tet-mS-HES1 (and thus mScarlet), a random subset of cells was followed independent of fluorescence. The mean percentage of non-dividing cells was calculated across biological replicates.

**Time Series Dip Size Quantification**

To quantify the magnitude of periodic dips in fluorescence intensity time-series, a custom MATLAB pipeline was developed. Peaks and troughs were detected using the *findpeaks* function, which returns both the amplitude and time index of features in the trace. Troughs were identified by applying the same function to the vertically inverted signal and taking the absolute values.

For circadian-level oscillations, traces were smoothed using *smoothdata* with a Gaussian filter and a moving window size of 10h, applied twice for enhanced smoothing. A minimum peak prominence of 0.5 was used to identify relevant features. For ultradian oscillations, the same process was applied using a 1.67h moving window and a minimum peak prominence of 1.5. All peak and trough amplitudes were extracted from the raw data at the corresponding time indices identified in the smoothed trace.

To assess dip magnitude, each peak was paired with its subsequent trough and dip amplitude (peak value – trough value) and fold-change (peak value / trough value) were calculated. In instances where a trace began with a trough, the preceding local maximum was designated as the initial peak, and vice versa for terminal peaks. Distributions of dip sizes were visualised in Rstudio using *ggplot2*.

**Periodicity Analysis Data Preprocessing**

Two-generation mother-daughter traces from proliferative DMSO control cells and uninterrupted palbociclib-arrested time-series were imported into MATLAB. All time-series were detrended using the *smoothdata* function. To identify circadian-level periodicity, a gaussian smoothing filter (with a window equal to the trace length which varied between 60-100h) was applied to yield a near-linear trend. This trend was then removed to even out intra and inter-cell intensity differences which may introduce bias to the data and obscure peak detection. For ultradian period quantification, a gaussian filter (10h moving window length) was applied and detrended. This was performed to remove periodicities greater than 10h, including the already-quantified circadian-level trend and reveal the nested ultradian oscillations.

**Autocorrelation Function**

Autocorrelation Function (ACF) quantifies the similarity of a signal with time-lagged versions of itself. Peaks in the ACF output correspond to repeated patterns, with the time between peaks providing a periodicity estimate. To assess robustness, a bootstrapping approach was used: each trace was randomly shuffled 100 times, and the ACF was computed on each permutation. Only ACF peaks in the original trace exceeding 0.5 standard deviations above the mean of the bootstrapped scores were retained. The mean time lag between these valid peaks was recorded as the periodicity estimate for that trace.

**Lomb-Scargle Periodogram**

Lomb-Scargle Periodogram (LSP) is a Fourier- and least-squares–based method optimised for non-uniformly sampled time-series data (VanderPlas et al., 2018). Periodograms were generated using the *plomb* function in MATLAB with the normalise flag to scale power by two times the variance of the time-series. Frequency–power spectra were interpolated across cells to generate a condition-wide average periodogram estimate. To better contextualise frequency analysis, the periodograms' x-axes were converted from frequency to period by taking the inverse of frequency. High-power, single-period peaks indicate strong oscillatory periodicity, while low, diffuse peaks reflect weak or aperiodic signals. LSP visualisations were generated using *ggplot2* in RStudio.

To statistically analyse LSP spectra from different signals, the dominant periodicity was extracted by converting the frequency at which the highest power is observed to a period value, on a cell-by-cell basis, by taking the reciprocal of the frequency. Mean power across circadian (20-30h) or ultradian periodicity range (4h-8h) was calculated by filtering for all frequencies which correspond to periodicities within the specified range, across all cells. Then the mean value of power was calculated at each frequency across every cell from all experiments. These mean power values within the specified range were then compared by boxplot in Rstudio using *ggplot2*.

**Kaplan-Meier survival curve analysis**

To investigate the effect of HES1 levels on survival in breast cancer, the Kaplan-Meier Plotter database (kmplot.com) was used (Posta *et al.*, 2025). Gene chip data for breast cancer was analysed, using the best probe for HES1 (203395_s_at) and automated cut-offs. Relapse-free survival was assessed in data from either Luminal A (PAM50) or ER+ (array) patients.

**Public RNA-seq Dataset Analysis**

Transcriptomics data in the form of bulk RNA sequencing was obtained from several sources including both MCF-7 cell line and breast cancer patient data to analyse HES1 expression, in support of our experimental observations. For all datasets, raw FASTQ files were downloaded from the NCBI Sequence Read Archive (SRA) using the SRA Toolkit (v3.0.7).

Data for palbociclib treatment in MCF-7s was obtained from Schmidt *et al.*, 2024 (GSE235350), and data on the effects of palbociclib discontinuation in MCF-7s was acquired from Armand *et al.*, 2025 (GSE279160). These studies were used to corroborate our finding that palbociclib treatment transiently increases HES1 levels. To characterise HES1 levels in breast cancer compared to normal mammary tissue, data from Li *et al.*, 2024 (GSE233242) was analysed, where RNA-seq was performed on paired biopsies of tumorous and normal breast epithelial cells from the same donor. Only sample pairs from Luminal A breast cancer donors were analysed. Additionally, HES1 expression was quantified from processed data provided by Frost, 2021, in which comparative analysis of 21 TCGA cancer types and their corresponding HPA normal tissue types were investigated. Data for HES1 (ENSG00000114315) expression from the Breast invasive carcinoma (BRCA) samples and normal breast tissue were extracted and graphically shown. This data corresponds to mean expression values in fragments per kilobase mapped reads (FPKM) from bulk RNA-seq for primary tumors provided by TCGA and normal breast tissues provided by the Human Protein Atlas (HPA).

Raw sequencing reads were quality-filtered and adapter-trimmed using Trimmomatic v0.36 (Bolger et al., 2014) with the following parameters: ILLUMINACLIP:<adapter file>:2:15:5 LEADING:3 TRAILING:3 SLIDINGWINDOW:4:20 MINLEN:20. Quality control was performed using FastQC v0.12.1 after trimming, confirming adapter removal and adequate read quality. Transcript abundances were quantified using Kallisto v0.51.1 (Andrews, 2010; Bray et al., 2016). A human index was constructed using the Ensembl GRCh38.110 cDNA reference and GTF annotation. Pseudo-alignment and abundance estimation were performed using default parameters for single and paired end samples, depending on the source data. Transcript per million (TPM) quantification was achieved as well as count matrices for all samples. To quantify the expression of HES1 across processed RNA-seq datasets, expression values corresponding to the canonical HES1 transcript (ENST00000232424.4) were extracted from Kallisto output and used for plotting. Differences between groups within each study were assessed using Welch's two sample t-test if sample numbers were greater than 3, performed in R (v4.5.0).

**Quantification and statistical analysis**

Statistical testing was performed in Rstudio v.4.4.1 using the '*stats'* and '*ggsignif'* libraries. Data was tested for normality using Shapiro-Wilk to select the appropriate test and significance. Statistical testing information, including significance values and sample size, is provided in figure legends.

**Supplementary Tables**

| **Primer pairs for genotyping mScarlet-I insertion following CRISPR** | |
|---|---|
| FP1 ⇆ RP1 | 5'-CCTTGCTCCGAAAAACCTGC-3' |
| | 5'-TGGACAATGCCTCCCAATCC-3' |
| FP2 ⇆ RP2 | 5'-AGCGCTACTGATCACCAAGT-3' |
| | 5'-CCATCCCGGGGTTTCAAGAC-3' |
| **Primer pairs for qPCR** | |
| HES1 | 5'-AAGGCGGACATTCTGGAAAT-3' |
| | 5'-GTCACCTCGTTCATGCACTC-3' |
| CDKN1A (p21) | 5'-GACTCTCAGGGTCGAAAACG-3' |
| | 5'-AAGATGTAGAGCGGGCCTTT-3' |
| GAPDH | 5'-CACCGTCAAGGCTGAGAACG -3' |
| | 5'-GCCCCACTTGATTTTGGAGG-3' |
| ActinB | 5'-GAAAATCTGGCACCACACCT-3' |
| | 5'-TAGCACAGCCTGGATAGCAA-3' |
| Rpl7 | 5'- ACAAGCGTGGTTATGGCAAA-3' |
| | 5'-CTCATGAATCAAATCCTCCATGCA-3' |

**Table S1. List of primers used for qPCR and to genotype the mS-HES1 SUM149 CRISPR cell line.**

### Supplementary Figures

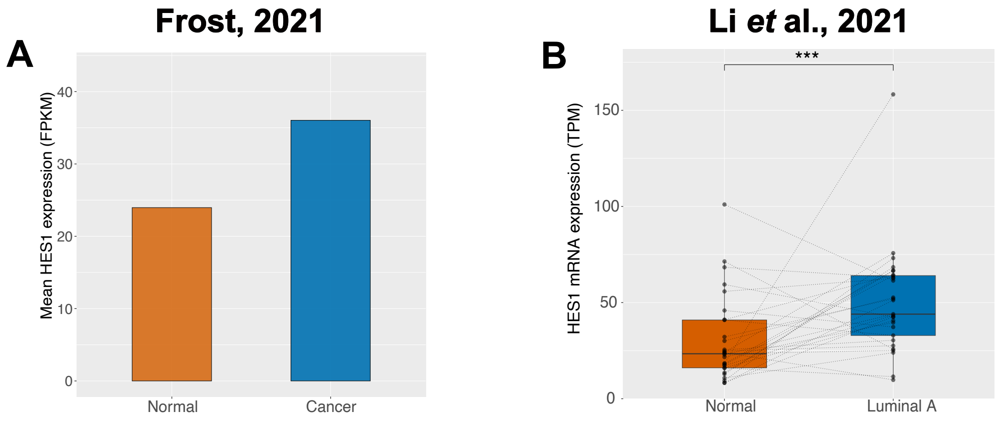


**Figure S1. HES1 is consistently upregulated in patient breast cancer samples compared to normal mammary epithelial tissue.**

(A) Mean expression values of HES1 in fragments per kilobase mapped reads (FPKM) in normal breast tissue versus cancer samples from RNA-seq. Processed data for normal breast cancer tissue (HPA) and breast cancer primary tumours (TCGA) was obtained from Frost, 2021.

(B) HES1 expression values, in transcripts per million (TPM), from RNA-seq of 29 paired primary Luminal A breast cancer cell and normal mammary epithelial cell samples from the same donor. Dotted lines indicate sample pairings. Paired Wilcoxon test ***p < 0.001.

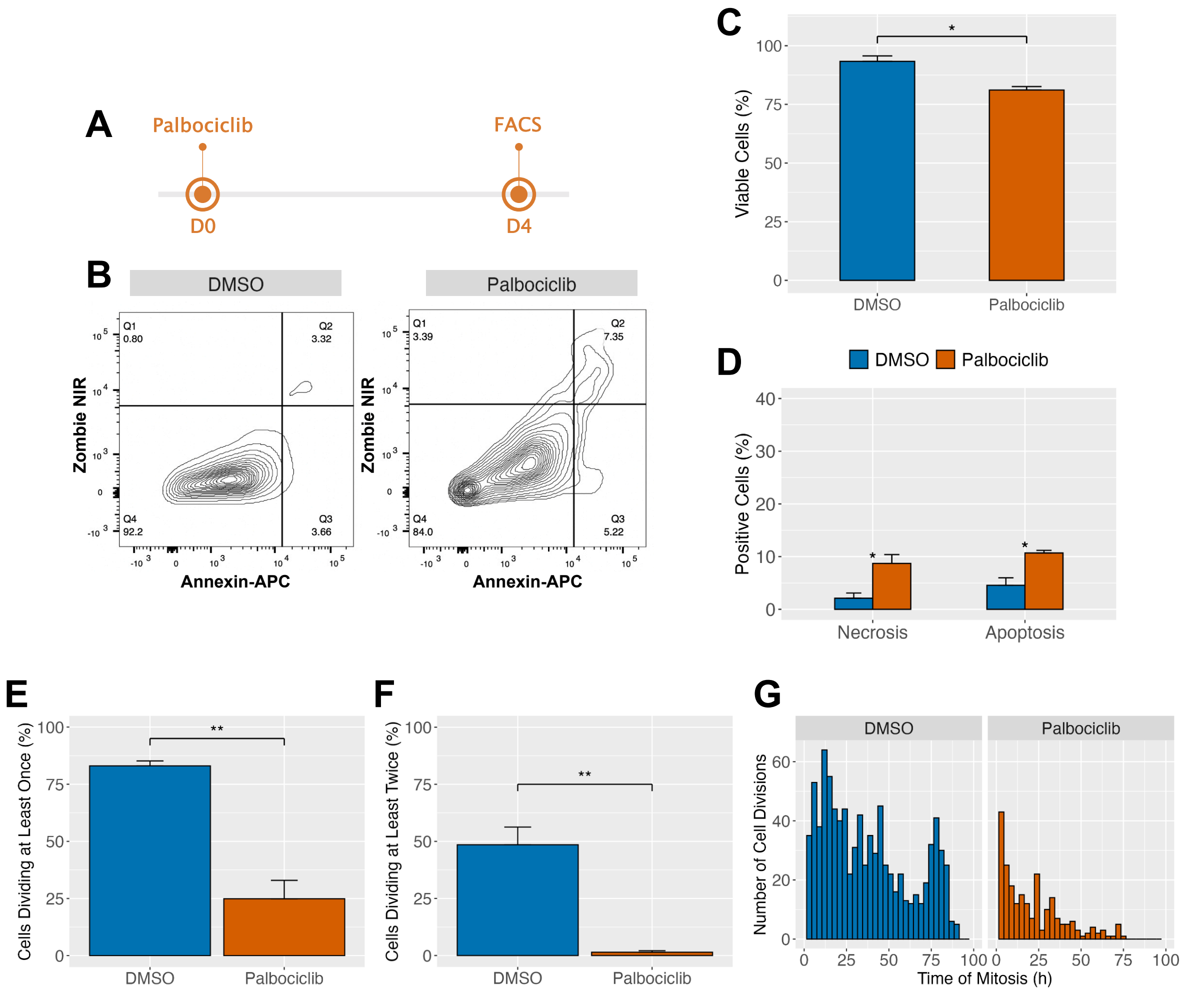


**Figure S2. Palbociclib impedes proliferation in the majority of cells and induces a small amount of cell death.**

(A) Timeline of (B-D), MCF-7 cells were treated with palbociclib or DMSO for 4 days then analysed by FACS.

(B) Representative examples of flow cytometry analysis of Annexin-APC and Zombie NIR viability dye staining.

(C, D) Quantification of the proportion of (C) viable cells, derived from Q4 in (B), and the proportion of (D) necrotic (Q1) and apoptotic (Q2 + Q3) cells. *N* = 3 experiments; *n* = 30,000 events. Error bars indicate SD. ANOVA with Tukey's post-hoc test. *p < 0.05.

(E, F) MCF-7 expressing endogenous mVenus-HES1 as a nuclear marker were treated with palbociclib, or DMSO as a vehicle-control, for 3 days and then continuously live-imaged. Divisions were defined as in (Fig. 1E). Bar charts showing the proportion of founder cells which divided at least once (E) or at least twice (F). *N* = 3 experiments; *n* = 526 (DMSO) or 2212 (palbociclib) founder cells. Paired t-test: **p < 0.01.

(G) Histogram shows the timing of all divisions across a field-of-view in each experimental condition.

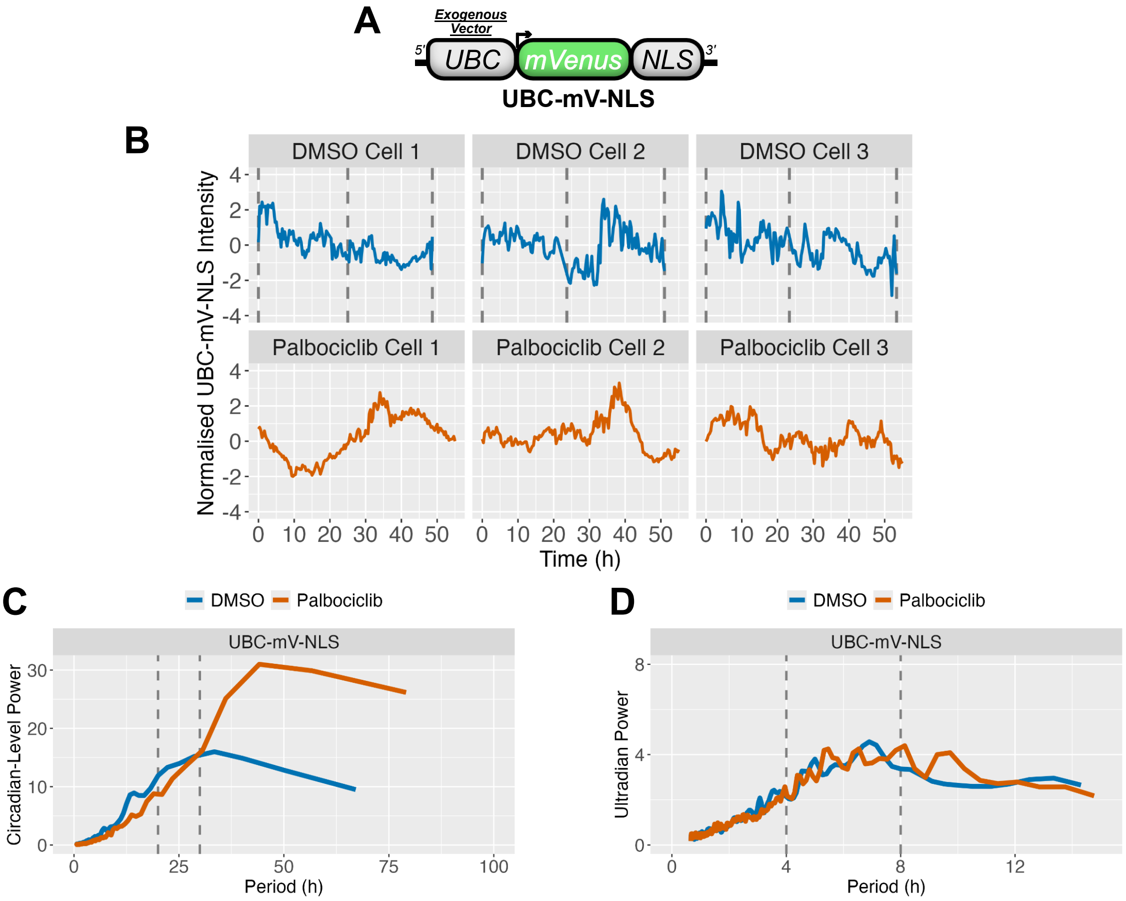


**Figure S3. A control UBC-mV-NLS reporter exhibits periodicity distinct from mV-HES1.**

(A) UBC-mV-NLS cells were continuously live-imaged in either proliferative (DMSO control) or arrested (palbociclib) conditions. Representative time-series from each group are shown. Time-series are Z-score normalised in order to control for inter-cell differences in levels. Grey lines indicate mitoses.

(B) LSP power spectra of time-series from proliferative (DMSO) or arrested (palbociclib) UBC-mV-NLS cells. Time-series were detrended to isolate circadian-level trends. Each spectrum represents the mean of all single-cell spectra from each group. A small broad peak is seen in DMSO cells indicating a lack of strong consensus periodicity. Palbociclib cells demonstrated a high power value but this did not form a peak and corresponded to period values approaching the length of the time-series, suggesting this is likely artefactual (VanderPlas, 2018; Hubner *et al.*, 2022; Casini *et al.*, 2024). *N* = 3 experiments, *n* = 90 cells per group.

(C) As in (B) but with time-series detrended to isolate ultradian oscillations (defined as 4–8h periods).

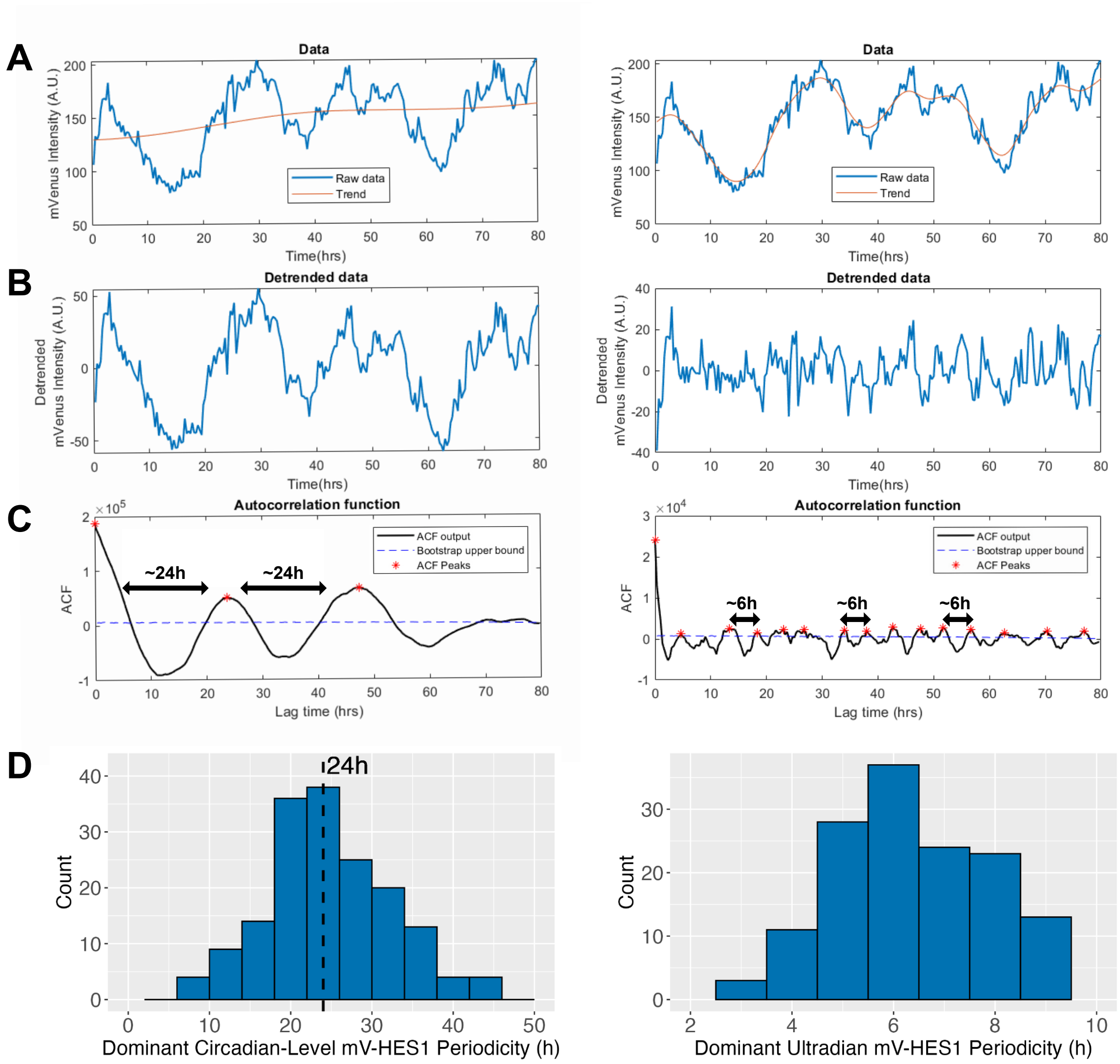


**Figure S4. Autocorrelation Function (ACF) pipeline to estimate circadian-level and ultradian HES1 periodicities.**

(A) Representative single-cell time-series of a proliferative mV-HES1 cell over two consecutive cell divisions. The same cell is shown alongside a linear trend (left) or a circadian-level trend (right).

(B) The trends identified in (A) were removed to isolate circadian-level (left) and ultradian traces (right).

(C) Autocorrelation function (ACF) applied to the detrended traces in (B), revealing recurrent peaks due to self-similarity. Peaks surpassing a bootstrap-derived threshold were classified as significant oscillatory peaks and are marked with red stars.

(D) Periodicity was estimated for each single cell by averaging the time intervals between ACF peaks. Histograms show the distribution of circadian-level (left) and ultradian (right) periodicities across the population. *N* = 3 independent experiments; *n* = 90 cells.

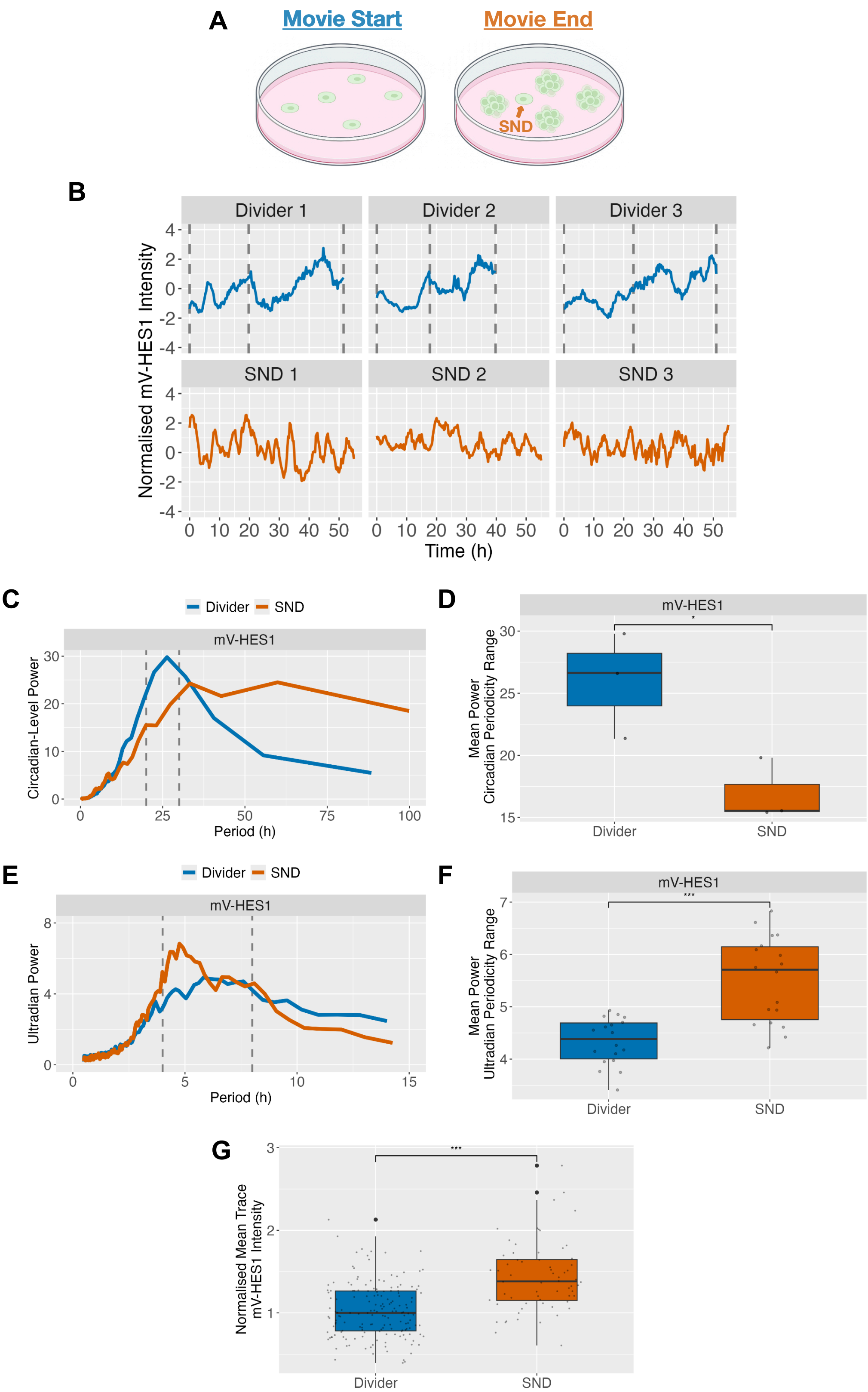


**Figure S5.** Unperturbed spontaneously non-dividing cells exhibit similar HES1 periodicity alterations as seen in palbociclib-arrested cells.

(A) Schematic illustrating the definition of spontaneous non-dividers (SNDs), a subset of mV-HES1 cells that, under proliferative conditions, spontaneously failed to divide for at least 60h (the average length of two generations in cycling cells) during the imaging time-course.

(B) Representative Z-score normalised HES1 intensity time-series from Divider and SND mV-HES1 cells. For Dividers, the x-axis represents time since the first mitosis. For SNDs, time is plotted from the beginning of the movie, as no mitosis occurred. Grey lines indicate mitoses.

(C) LSP power spectra of detrended time-series from Divider and SND mV-HES1 cells, isolating circadian-level oscillations. Spectra represent the mean of all single-cell power spectra within each group. $N = 3$ experiments, $n = 164$ (Dividers), 64 (SND).

(D) Comparison of circadian-range (20–30h) oscillatory power between dividing and SND cells, based on spectra in (C). Each data point represents mean power at a specific frequency. $N = 3$ experiments, $n = 164$ (Dividers), 64 (SND). Statistical significance was assessed using an independent t-test. *$p < 0.05$.

(E, F) As in (C) and (D), but with detrending to isolate ultradian oscillations (4–8h range). ***$p < 0.001$.

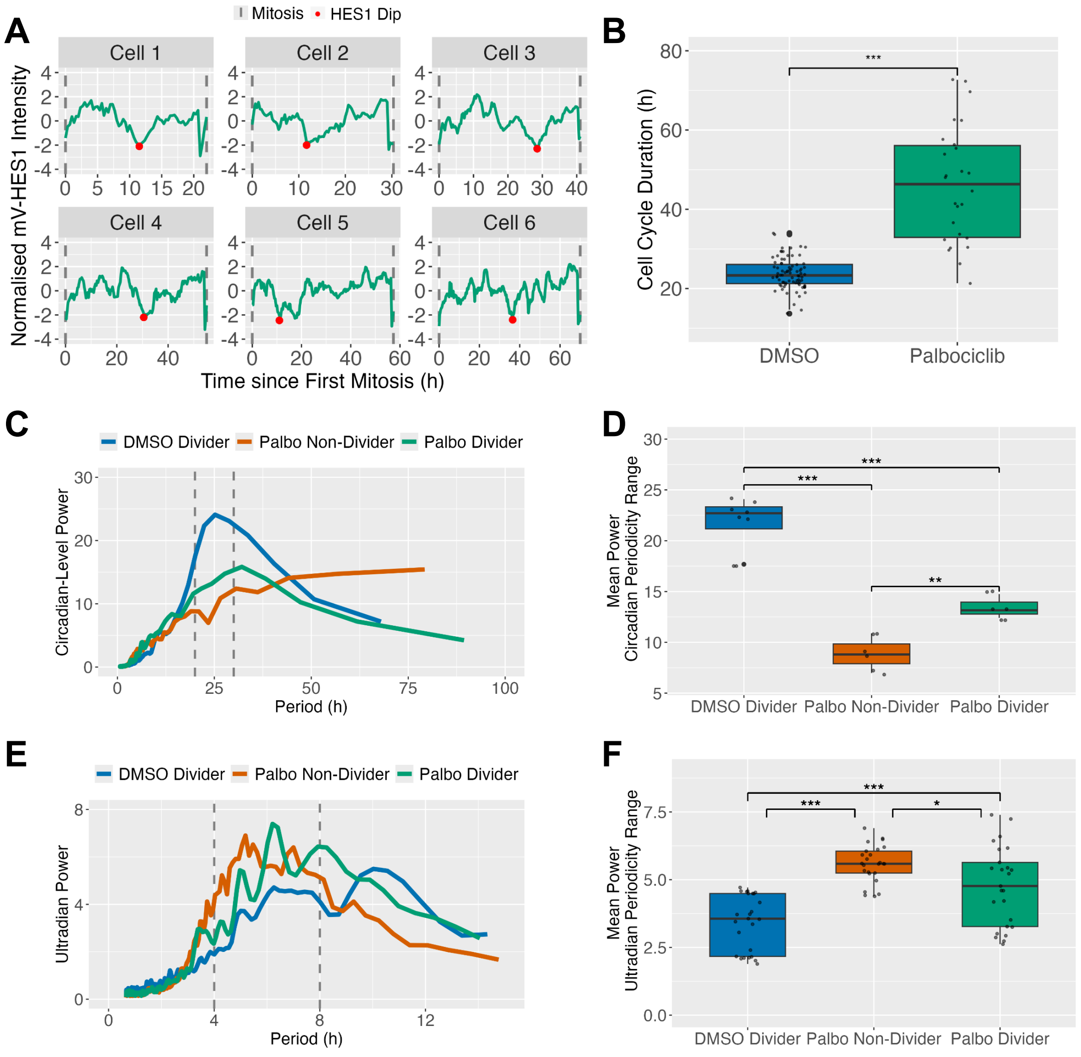


**Figure S6.** Cells which continue to divide in palbociclib exhibit longer cell cycles and intermediate periodicity.

(A) Representative Z-score normalised mV-HES1 intensity time-series of cells which escaped arrest and completed a full cell cycle during day 3-7 of palbociclib treatment. Cells of increasing cell cycle duration are shown from Cell 1 to Cell 6.

(B) Cell cycle duration of cells in proliferative conditions (DMSO) and those which escaped palbociclib treatment. $N = 3$ experiments; $n = 80$ (DMSO) or 26 (palbociclib dividers) cells. Independent t-test: ***p $< 0.001$.

(C) LSP power spectra of detrended time-series isolating circadian-level oscillations from palbociclib divider cells overlaid with DMSO dividers and palbociclib non-dividers. Spectra represent the mean of all single-cell power spectra within each group. For (C-F), *N* = 3 experiments, *n* = 90 (DMSO dividers and palbociclib non-dividers), 26 (palbociclib dividers).

(D) Comparison of circadian-range (20–30h) oscillatory power between groups, based on spectra in (C). Each data point represents mean power at a specific frequency. ANOVA with Tukey's post-hoc test. ***p < 0.001, **p < 0.01.

(E, F) As in (C) and (D), but with detrending to isolate ultradian oscillations (4–8h range). *p < 0.05.

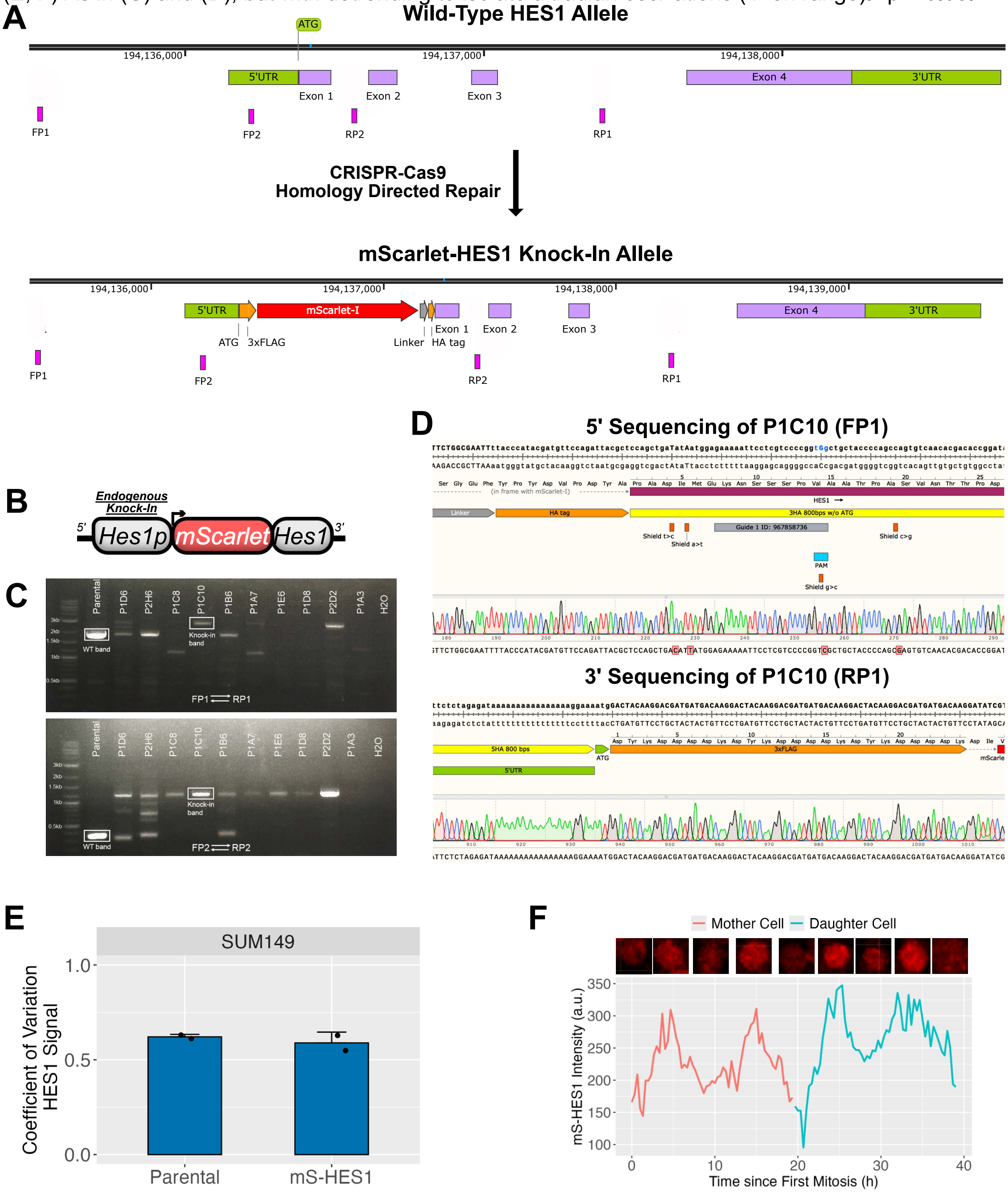


**Figure S7. Genotyping and validation of a CRISPR knock-in mediated endogenous HES1 fusion-reporter in the triple-negative breast cancer line SUM149.**

(A) Schematic of the HES1 genomic locus before (top) and after (bottom) endogenous knock-in of mScarlet at the N-terminus by CRISPR-Cas9 mediated homology directed repair (HDR). The binding sites of several genotyping primers are shown (FP1, RP1, FP2, RP2).

(B) Schematic of the endogenous HES1 promoter-driven mScarlet-HES1 fusion reporter (mS-HES1).

(C) PCR genotyping of a multiple clonal cell lines using the FP1/RP1 (top) or FP2/RP2 (bottom primers). Successful homozygous knock-in band in the selected mS-HES1 cell line (clone P1C10) is highlighted, alongside the wild-type endogenous band in parental SUM149s.

(D) Sequencing results from TOPO-cloned alleles of P1C10 at the 5' (top) and 3' (bottom) ends. Silent shield mutations near the PAM site which were intentionally designed to prevent cutting during the CRISPR-Cas9 reaction are shown. No other mutations were found.

(E) Immunostaining for HES1 was performed to then analyse the coefficient of variation (CV) of HES1 in parental SUM149s and the mS-HES1 knock-in (P1C10), serving as a readout of population-level HES1 heterogeneity which is used as a proxy for asynchronous single-cell fluctuations. *N = 2* biological replicates; *n* = ~3000 cells per group.

(F) Representative single-cell time-series of mS-HES1 fluorescence intensity in a mother-daughter pair, tracked for two full cell cycle generations from mitosis-to-mitosis. Corresponding fluorescence images from this cell are shown above the time-series.

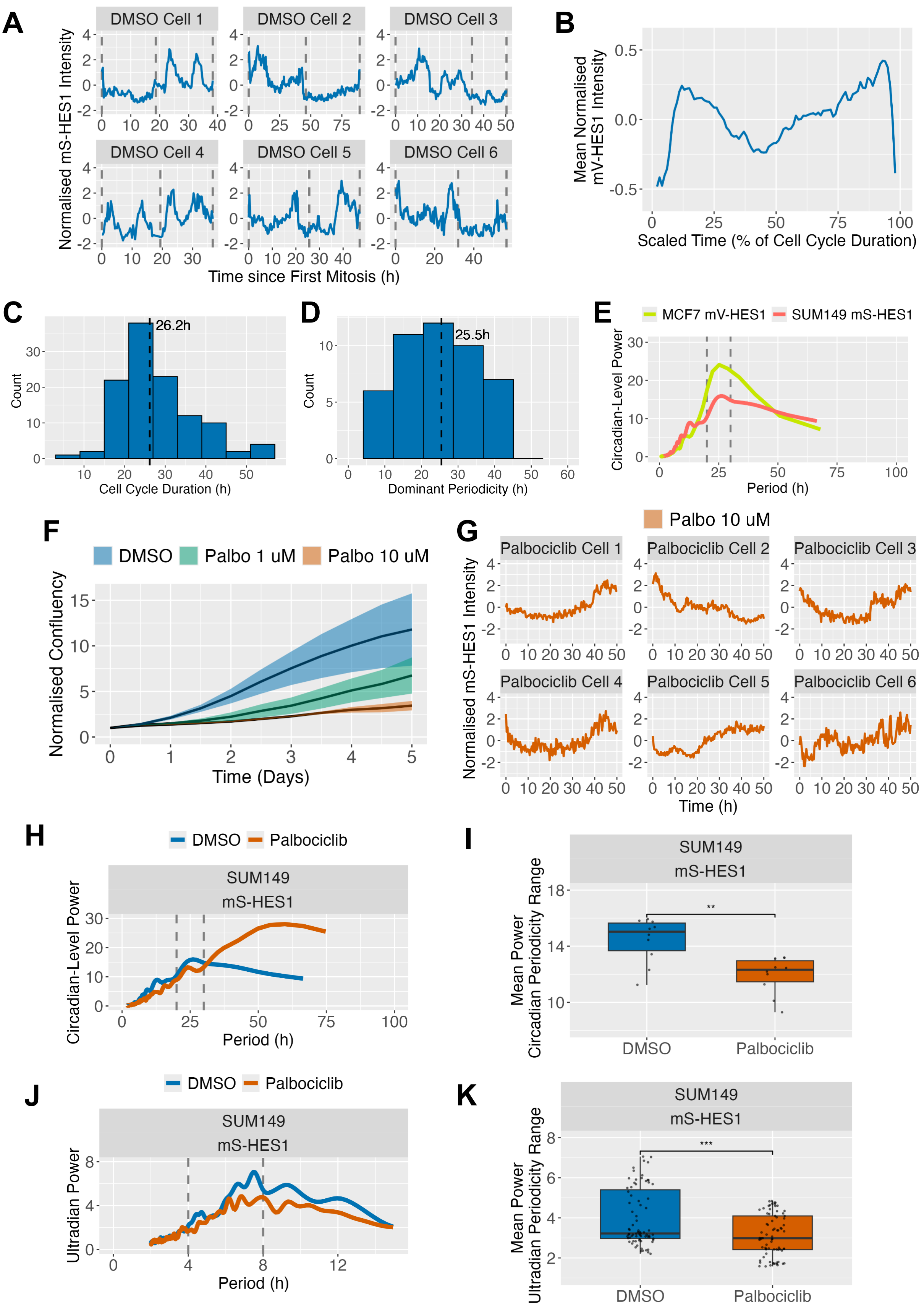


**Figure S8. SUM149 cells also exhibit circadian-level HES1 oscillations which are depleted during palbociclib-mediated arrest.**

(A) Representative single-cell Z-score normalised mS-HES1 intensity time-series of proliferative SUM149 cells over 2 cell cycle generations. Grey lines indicate mitoses.

(B) Mean trace of (A), with time scaled a percentage of each cell's cell cycle duration.

(C) Histogram to show the distribution of cell cycle durations of proliferative mS-HES1 SUM149 cells. Black line indicates median cell cycle duration.

(D) Histogram to show estimated circadian-level periodicities in proliferative mS-HES1 SUM149 cells derived from Lomb-Scargle periodogram (LSP) approach. *N* = 3 independent experiments *;* *n* = 50 cells.

(E) LSP power spectra of detrended time-series isolating circadian-level oscillations from proliferative mV-HES1 MCF-7s and mS-HES1 SUM149s. Spectra represent the mean of all single-cell power spectra within each group.

(F) Population-level growth curves of SUM149 cells treated with DMSO and 1 μM or 10 μM palbociclib. Confluency was normalised to baseline values at the start of the time course. Lines represent the mean of *N* = 3 biological replicates; shaded areas show standard deviation (SD).

(G) Representative Z-score normalised mS-HES1 intensity time-series of palbociclib-arrested SUM149 cells.

(H) Circadian-level LSP power spectra comparing mS-HES1 SUM149s treated with DMSO or palbociclib. Spectra represent the mean of all single-cell power spectra within each group. *N* = 3 experiments, *n* = 50-55 cells.

(I) Comparison of circadian-range (20–30h) oscillatory power between groups, based on spectra in (E). Each data point represents mean power at a specific frequency. ANOVA with Tukey's post-hoc test. **p < 0.01.

(J, K) As in (H) and (I), but with detrending to isolate ultradian oscillations (4–8h range). ***p < 0.001.

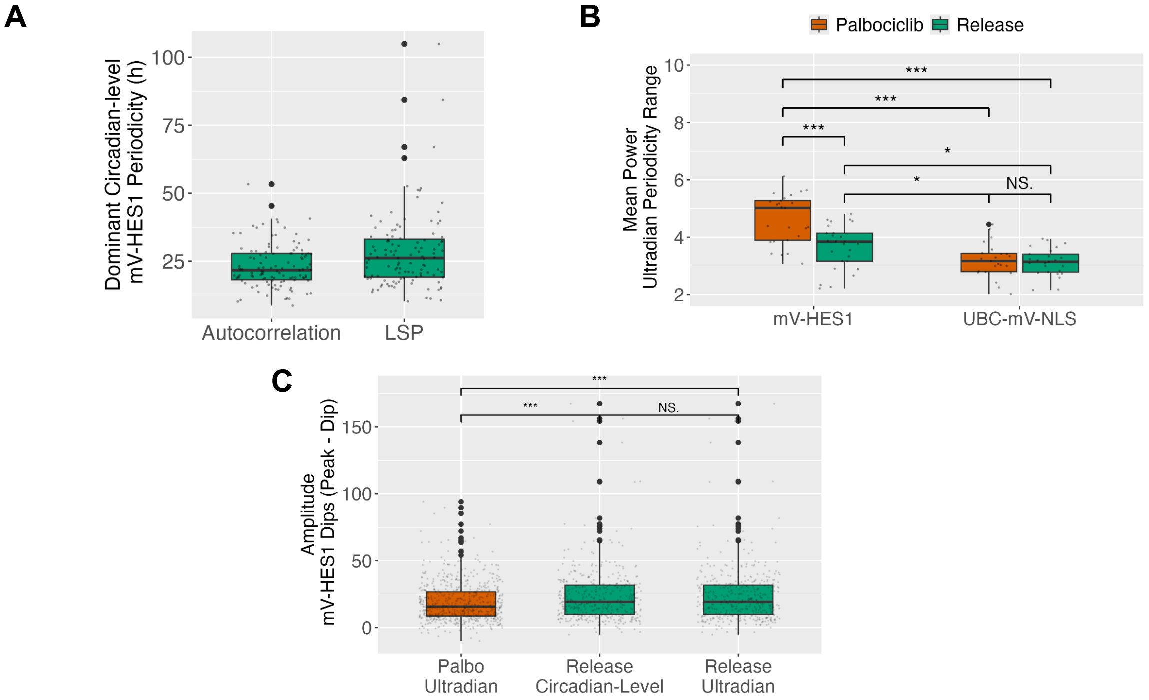


**Figure S9. HES1 ultradian periodicity is reduced and dip amplitude increases as cells re-enter the cell cycle, reversing changes seen during arrest.**

(A) Estimation of HES1 circadian-level periodicities in mV-HES1 cells following release from palbociclib-mediated arrest. Periodicities were assessed using two independent methods: autocorrelation and Lomb-Scargle periodogram (LSP). $n$ = 115 cells pooled across $N$ = 3 experiments.

(B) Comparison of ultradian-range (4–8h) oscillatory power between released and palbociclib-treated cells. Each point represents mean power at a specific ultradian frequency averaged across all cells and replicates. Statistical significance was assessed using ANOVA with Tukey's post-hoc test. All significance comparisons shown. ***p < 0.001, *p < 0.05.

(C, D) Fold-change (C) and amplitude (D) of every peak-dip in HES1 intensity traces across all cells. Comparisons were made between released cells detrended for circadian-level periodicity, released cells detrended for ultradian periodicity and palbociclib-treated cells detrended for ultradian periodicity. $N$ = 3 experiments, $n$ = 115 (Release), 90 (Palbociclib). Statistical analysis was performed using ANOVA with Tukey's post-hoc test. ***p < 0.001.

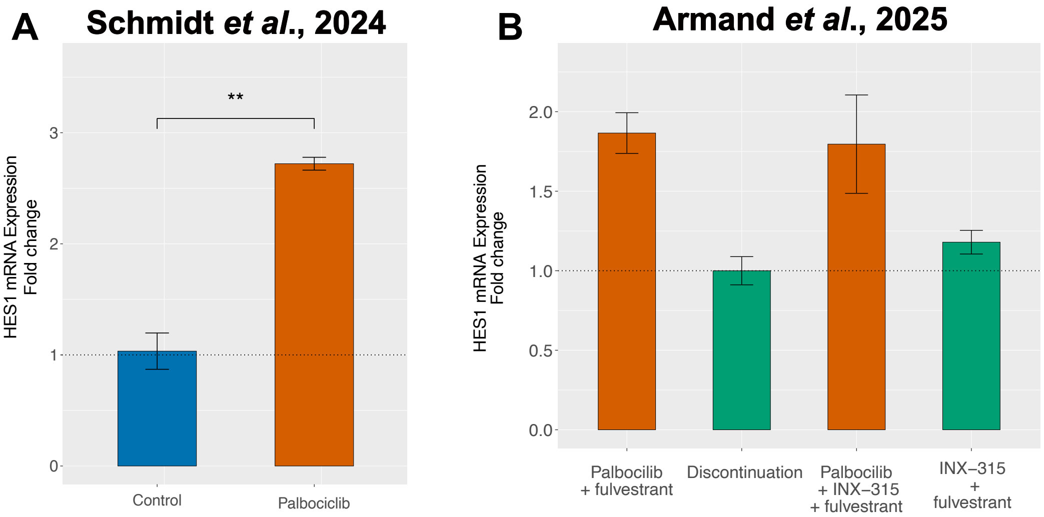


**Figure S10. HES1 levels are also reversibly elevated by palbociclib in independent studies.**

(A) HES1 expression values during palbociclib treatment in MCF-7 cells, measured by RNA-seq and expressed as fold change in TPM from the control sample. Analysed from publicly-available data (Schmidt *et al*., 2024). N = 3 biological replicates per sample. Paired t test **p < 0.01.

(B) HES1 expression data from publicly-available RNA-seq data (Armand *et al*., 2025), analysed as in (A). MCF-7 cells were treated with a combination of palbociclib and fulvestrant for 2 months which was then discontinued by switching to either DMSO or INX-315 (CDK2 inhibitor) and fulvestrant ± palbociclib for 2 weeks before harvesting samples for RNA-seq. N = 2 biological replicates per sample.

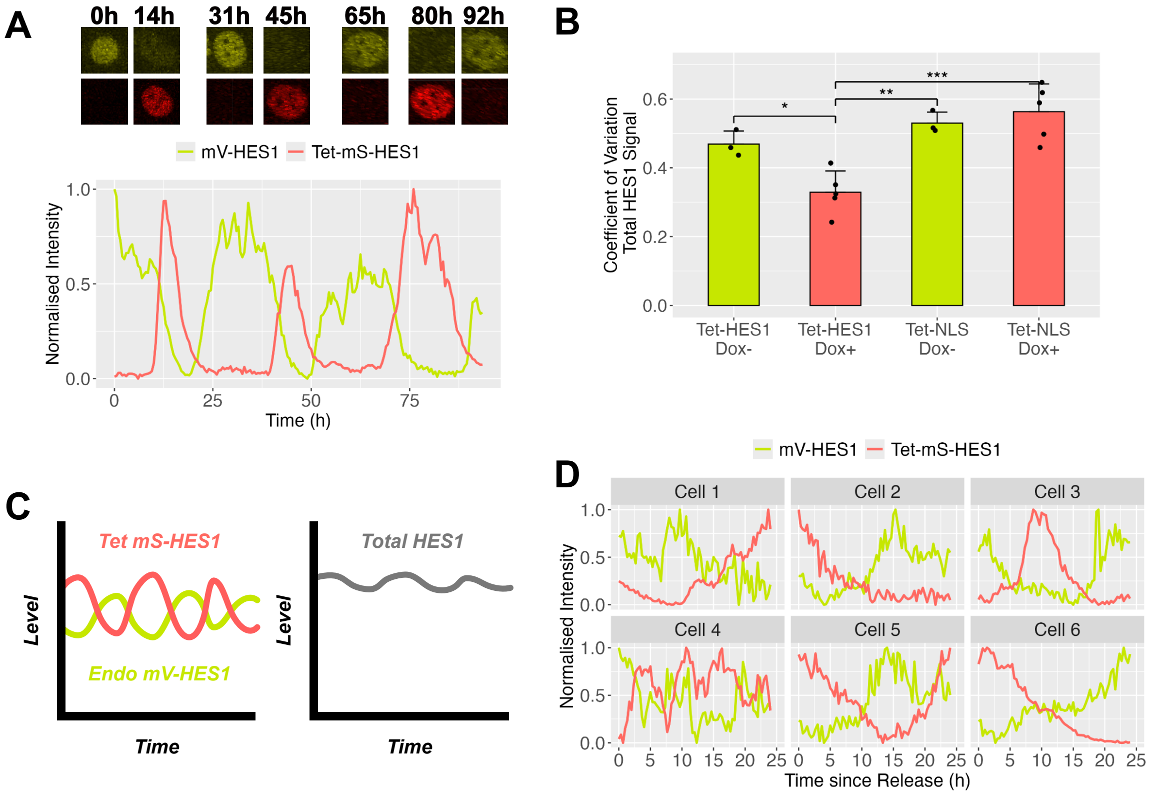


**Figure S11. Anti-phase Tet-mS-HES1 and endogenous mV-HES1 oscillations result in more sustained total HES1 expression.**

(A) Representative single-cell example of a mV-HES1 cell expressing the doxycycline-inducible Tet-mS-HES1 reporter and showing anti-phase oscillations between the two reporters. Nuclear images corresponding to the displayed mV-HES1 and Tet-mS-HES1 traces are overlaid above.

(B) mV-HES1 cells expressing either Tet-mS-HES1 or a control Tet-mS-NLS reporter were treated with doxycycline for 3 days, followed by total HES1 immunostaining to measure combined endogenous (mV-HES1) and exogenous (mS-HES1) protein levels. The coefficient of variation (CV) across single cells was used as a population-level proxy for asynchronous single-cell fluctuations. Tet-mS-HES1–positive cells showed reduced CV relative to control cells, consistent with more sustained total HES1 levels. *N = 3* biological replicates; *n = ~15,000* cells per group. Dox+ samples include two replicates treated with 5 ng/mL and three with 50 ng/mL doxycycline. ANOVA with Tukey's post-hoc test. ***p < 0.001, **p < 0.01, *p < 0.05.

(C) Schematic illustrating how anti-phase oscillations of exogenous Tet-mS-HES1 and endogenous mV-HES1 may amount to the observed reduction in variation, interpreted as less dynamic, more sustained total HES1 (grey line).

(D) Representative time-series from 24h re-entry assay in Fig. 6. Doxycycline-induced mV-HES1 Tet-mS-HES1 cells were arrested with palbociclib for 3 days, then released into proliferative media. Traces normalised as in (A).

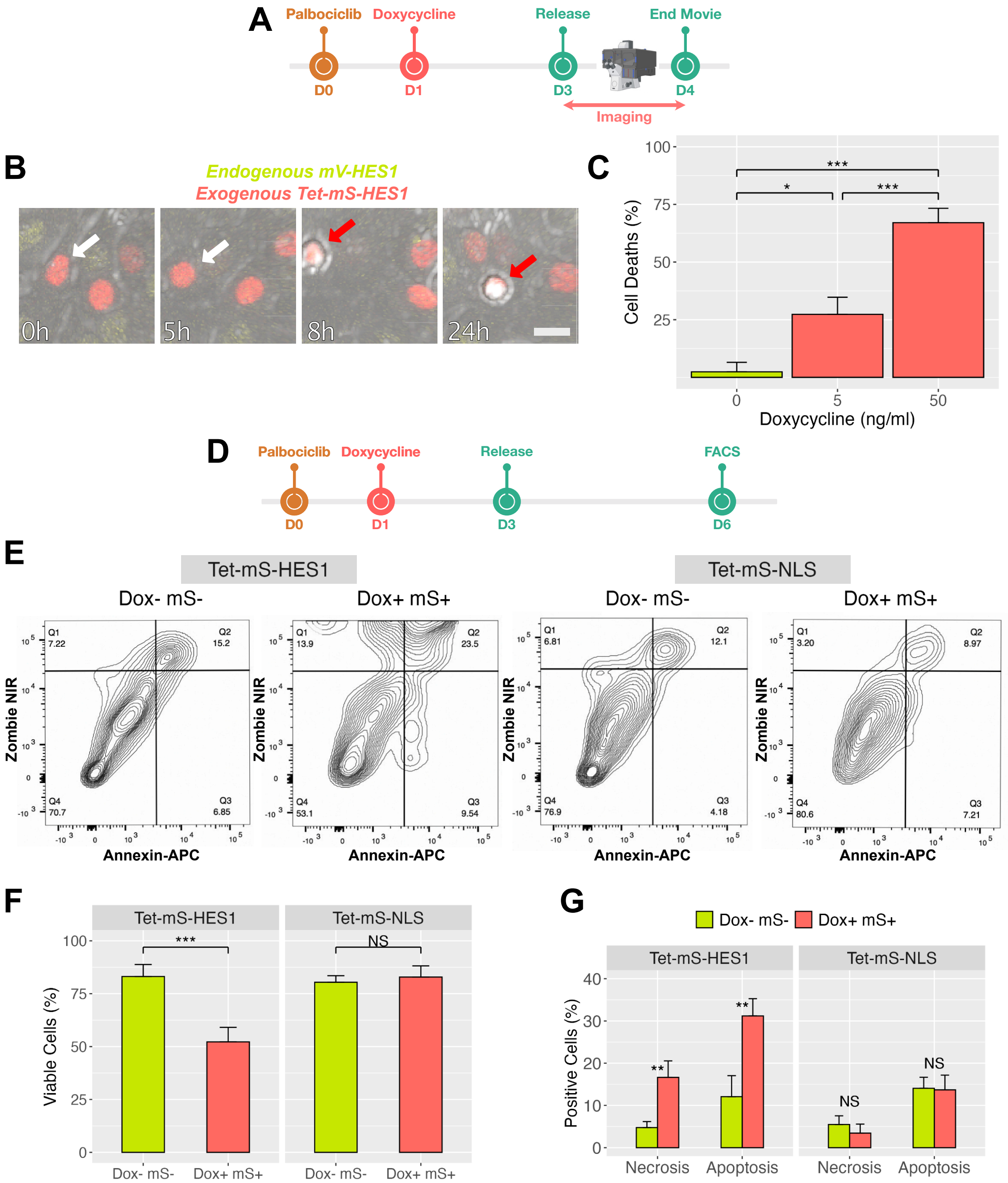


**Figure S12. Sustained HES1 induces cell death in cells released from palbociclib-mediated arrest.**

(A) Experimental timeline of (B-C). mV-HES1 ells containing the Tet-mS-HES1 cassette were released from palbociclib-mediated arrest ± doxycycline and continuously imaged for 24h.

(B) Cell deaths during this time-course were manually designated based on cell morphology (see methods). Example time-course of a Tet-mS-HES1 cell which undergoes cell death is shown. White and red arrows indicates the cell before and after death, respectively. Scale bar = 20µm.

(B) Quantification of the proportion of cell deaths observed during the 24h time-course in (A), in response to escalating concentrations of doxycycline. Bars show the mean proportion from $N$ = 3 experiments; $n$ = 186–199 cells per group. Error bars indicate SD. Independent t-test: ***$p < 0.001$, *$p < 0.05$.

(D) Experimental timeline of (E-G). Tet-mS-HES1 or control Tet-mS-NLS cells were released from palbociclib-mediated arrest ± doxycycline and were analysed by FACS 3 days later.

(E) Representative examples of flow cytometry analysis of Annexin-APC and Zombie NIR viability dye staining.

(F, G) Quantification of the proportion of (F) viable cells, derived from Q4 in (E), and the proportion of (G) necrotic (Q1) and apoptotic (Q2 + Q3) cells. $N$ = 3 experiments; $n$ = 30,000 events. Error bars indicate SD. ANOVA with Tukey's post-hoc test. ***p < 0.001, **p < 0.01.

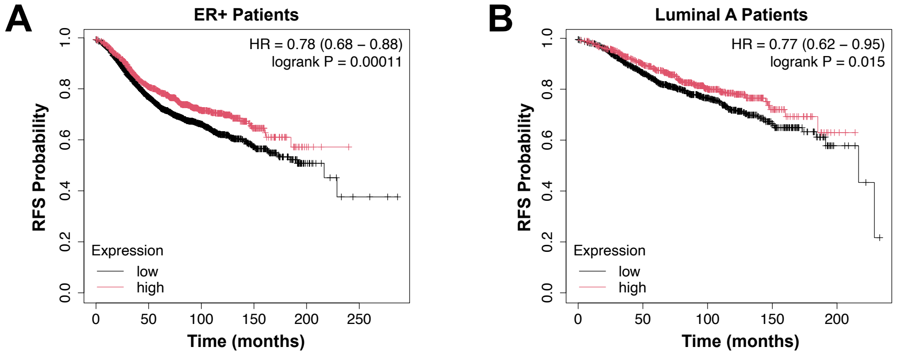


**Figure S13. High HES1 levels are associated with prolonged relapse-free survival in ER+ and Luminal A breast cancer patients.**

(A, B) Kaplan-Meier curves depicting relapse-free survival rates in (A) ER+ and (B) Luminal A breast cancer patients, based on high or low levels of HES1 expression. Data acquired from Posta *et al.*, 2025.

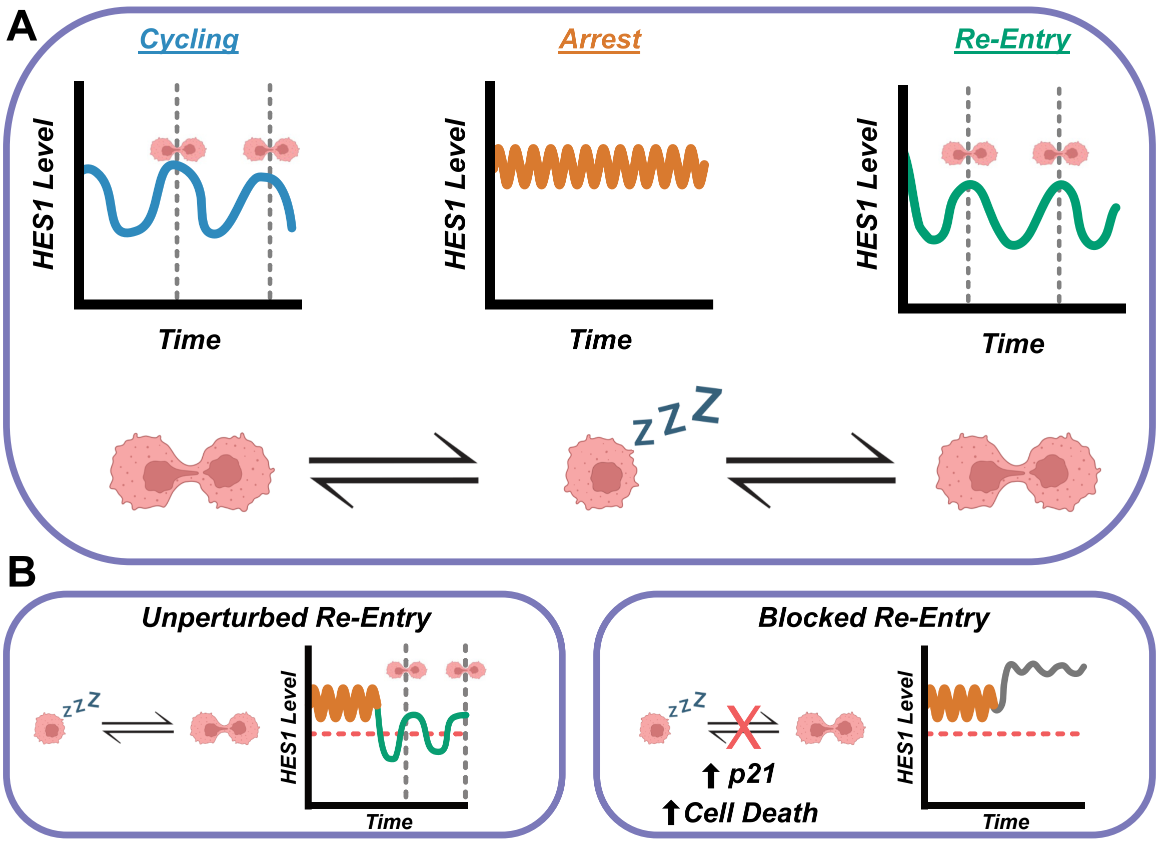


**Figure S14. Graphical Abstract.**

(A) HES1 exhibits 24h oscillations in cycling cells which are lost in arrested cells and replaced with less dynamic, high-frequency fluctuations. Upon cell cycle re-entry, a prominent dip is seen during the G1/S transition and 24h oscillations resume. Thus, transitions between distinct patterns of HES1 dynamics are observed during transitions between proliferation and arrest.

(B) Cell cycle re-entry can be perturbed by manipulating the dynamics of HES1. Inducing expression of elevated and less dynamic HES1 as cells attempt to re-enter the cell cycle, prevents the ordinarily-observed G1/S dip, upregulates the cell cycle inhibitor p21 and induces cell death. Therefore, manipulating HES1 blocks cell cycle re-entry and prevents population outgrowth.

### SI Movie Legends

**Movie S1. DMSO-treated control proliferative mV-HES1 cells.**

Movie depicts mV-HES1 MCF7 cells treated with DMSO for 5 days and then continuously imaged for 48h. Visible yellow signal is mV-HES1. Mitoses are visible when nuclear envelope breaks down and mV-HES1 signal diffuses into the cytoplasm, followed by cytokinesis.

**Movie S2. Palbociclib-treated arrested mV-HES1 cells.**

Movie depicts mV-HES1 MCF7 cells treated with palbociclib for 5 days and then continuously imaged for 48h. Visible yellow signal is mV-HES1. Majority of cells do not undergo cell division indicating cell cycle arrest.

**Movie S3. Released mV-HES1 cells.**

Movie depicts mV-HES1 MCF7 cells which were arrested with palbociclib for 3 days, prior release into proliferative media and then continuously imaged for 96h. Visible yellow signal is mV-HES1. Majority of cells exhibit cell division, signifying efficient cell cycle re-entry. A synchronous wave of re-entries are visible at 18-20h. Subsequent divisions are less globally synchronised.

**Movie S4. Induction of Tet-mS-HES1 impedes cell cycle re-entry.**

Movie depicts mV-HES1 MCF7 cells which also expressed the Tet-mS-HES1 cassette. Cells were arrested with palbociclib for 3 days, including doxycycline treatment after 24h. Then released into proliferative media, containing doxycycline and continuously imaged for 24h to assess cell cycle re-entry. For clarity, only Tet-mS-HES1 signal is shown. Cells which were positive for Tet-mS-HES1 at the time of release, and thus experienced experimentally sustained HES1 at the correct moment, are marked with a spot and tracked throughout. The overwhelming proportion of which (18/20, 90%, in this field of view) do not undergo cell division (see Fig. 6 for further quantification).
